# Analysis of genomic loci harboring 59,732 human-specific regulatory sequences reveals unique to human regulatory patterns associated with brain development

**DOI:** 10.1101/432625

**Authors:** Gennadi V. Glinsky

## Abstract

Extensive searches for genomic regions harboring various types of candidate human-specific regulatory sequences (HSRS) identified thousands’ HSRS using high-resolution next-generation sequencing technologies and methodologically diverse comparative analyses of human and non-human primates’ reference genomes. Here, a comprehensive catalogue of 59,732 genomic loci harboring candidate HSRS has been assembled to facilitate the systematic analyses of genomic sequences that were either inherited from extinct common ancestors (ECAs) or created de novo in human genomes. Present analyses identified thousands of HSRS that appear inherited from ECAs yet absent in genomes of our closest evolutionary relatives, Chimpanzee and Bonobo, presumably due to the incomplete lineage sorting and/or species-specific loss or regulatory DNA. This pattern is particularly prominent for HSRS that have been putatively associated with human-specific (HS) gene expression changes in cerebral organoid models. Significant fractions of retrotransposon-derived loci transcriptionally-active in human dorsolateral prefrontal cortex (DLPFC) are highly conserved in genomes of Gorilla, Orangutan, Gibbon, and Rhesus (1,688; 1,371; 1,148; and 1,045 loci, respectively), yet they are absent in genomes of both Chimpanzee and Bonobo. A prominent majority of regions harboring HS mutations associated with HS expression changes during brain development is highly conserved in Chimpanzee, Bonobo, and Gorilla genomes. Among non-human primates (NHP), dominant fractions of HSRS associated with HS gene expression in both excitatory neurons (347 loci; 67%) and radial glia (683 loci; 72%) are highly conserved in the Gorilla genome. Analysis of 4,433 genes encoding virus-interacting proteins (VIPs) revealed that 95.9% of human VIPs are components of HS regulatory networks that appear to operate in distinct types of human cells from preimplantation embryos to adult DLPFC. Present analyses demonstrate that Modern Humans captured unique combinations of regulatory sequences, divergent subsets of which are highly conserved in distinct species of six NHP separated by 30 million years of evolution. Concurrently, this unique-to-human mosaic of genomic regulatory patterns inherited from ECAs was supplemented with 12,486 created de novo HSRS. Present analyses of HSRS support the model of complex continuous speciation process during evolution of the human lineage that is not likely to occur as an instantaneous event. Genes encoding VIPs may represent a principal genomic target of HS regulatory networks, thus affecting a functionally diverse spectrum of biological processes controlled by VIP-containing liquid-liquid phase separated condensates.

## Introduction

Recent advances enabled by the analyses of individual genomes of Great Apes using high-resolution sequencing technologies and methodologically diverse comparative analyses of human and non-human primates’ reference genomes significantly enhanced our understanding of human-specific structural genomic variations of potential regulatory and functional significance (Locke et al., 2005; Chimpanzee Sequencing and Analysis Consortium, 2005; McLean et al., 2011; Prüfer et al., 2012; Shulha et al., 2012; Konopka et al., 2012; Scally et al., 2012; Capra et al., 2013; Marchetto et al., 2013; Marnetto et al., 2014; Prescott et al., 2015; Gittelman et al. 2015; Glinsky et al., 2015-2018; Dong et al., 2016; Sousa et al., 2017; Dennis et al., 2017; Kronenberg et al., 2018; Guffanti et al., 2018). Collectively, these studies markedly expanded the compendium of candidate human-specific regulatory sequences (HSRS), which currently comprises nearly sixty thousand genomic loci aligned to the most recent release of the human reference genome (Tables 1-4; Supplemental Tables S1-S12). This remarkable progress highlights a multitude of significant contemporary challenges, the centerpiece of which is a need to compile a comprehensive catalog of HSRS in order to identify the high-priority panel of genetic targets for stringent functional validation experiments. The selected high-priority genetic panel of human phenotypic divergence would represent the elite set of HSRS, which will be chosen based on the expectation of high-likelihood of biologically-significant species-specific effects on phenotypes that would be revealed during in-depth structural-functional explorations of their impact on development of human-specific traits.

One of essential steps toward addressing this problem is to gain insights into evolutionary origins of genomic regions harboring HSRS. In this contribution, an up-to-date catalog of 59,732 candidates human-specific regulatory loci has been assembled and their conservation patterns in genomes of five non-human Great Apes (Chimpanzee, Bonobo, Gorilla, Orangutan, and Gibbon; Tables 1-4; Supplemental Table S1-S12) have been analyzed. Systematic comparisons of genomic coordinates of HSRS and unique to African Great Apes insertions of the PtERV1 retrovirus-derived sequences were carried-out by performing comprehensive genome-wide proximity placement analyses. Diverse patterns of sequence conservation of different classes of HSRS were observed, reflecting quantitatively distinct profiles of inheritance from extinct common ancestors (ECAs) of the human lineage and each of the five species of non-human Great Apes. One of the prevalent modes of sequence conservation is represented by the bypassing pattern of evolutionary inheritance, which is exemplified by thousands of HSRS that appear inherited from ECAs yet bypassed genomes of our closest evolutionary relatives, Chimpanzee and Bonobo. The bypassing pattern of evolutionary inheritance seems particularly prominent for candidate HSRS putatively associated with development and functions of human brain.

## Results

### Mosaicism of evolutionary origins of genomic loci harboring various classes of human-specific mutations

Recent experiments identified 24,151 genomic regions harboring various classes of human-specific mutations identified based on the comparative analyses of genomes of Modern Humans and non-human Great Apes (Kronenber et al., 2018). It was of interest to analyze the sequence conservation patterns of these regions in genomes of six non-human primates (NHP), including five non-human Great Apes (Chimpanzee, Bonobo, Gorilla, Orangutan, and Gibbon) and Rhesus Macaque (Table 1). In these analyses, genomic sequences that manifested at least 95% of sequence conservations during the direct and reciprocal conversions from/to reference genomes of Modern Humans (hg38) and corresponding NHP species were considered highly-conserved. Within the context of definition of evolutionary origins of genomic regions harboring human-specific mutations, one of the main motivations was the inference that this analytical effort would identify highly-conserved DNA sequences that were inherited by the Modern Humans’ lineage from ECAs.

**Table 1.**
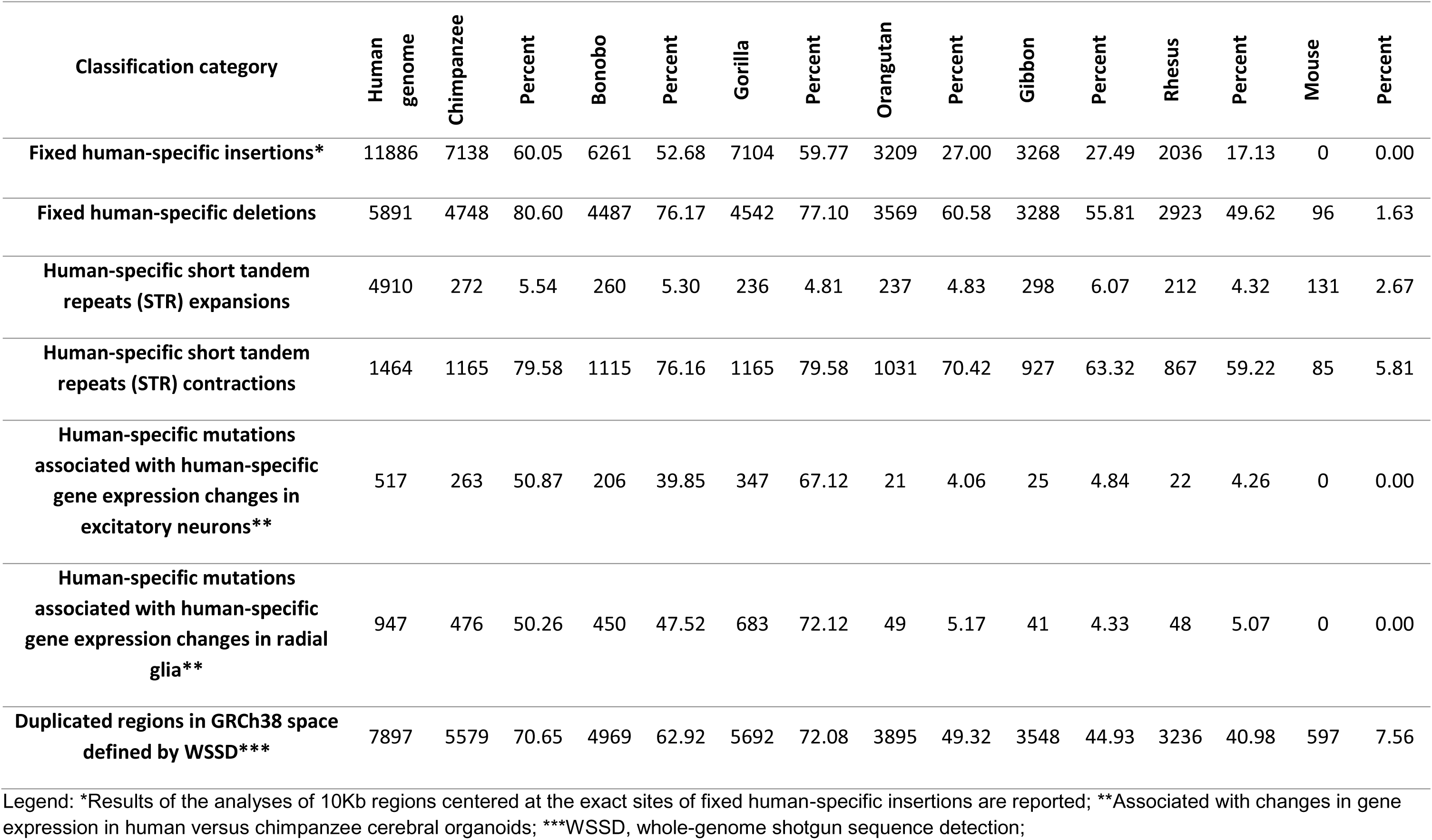
Mosaicism of evolutionary origins of genomic loci harboring various classes of human-specific mutations: Human-specific mutations target highly conserved sequences mapped to reference genomes of multiple species of non-human primates (95% sequence identity thresholds during both direct and reciprocal conversions).

| Classification category | Human genome | Chimpanzee | Percent | Bonobo | Percent | Gorilla | Percent | Orangutan | Percent | Gibbon | Percent | Rhesus | Percent | Mouse | Percent |
| --- | --- | --- | --- | --- | --- | --- | --- | --- | --- | --- | --- | --- | --- | --- | --- |
| **Fixed human-specific insertions\*** | 11886 | 7138 | 60.05 | 6261 | 52.68 | 7104 | 59.77 | 3209 | 27.00 | 3268 | 27.49 | 2036 | 17.13 | 0 | 0.00 |
| **Fixed human-specific deletions** | 5891 | 4748 | 80.60 | 4487 | 76.17 | 4542 | 77.10 | 3569 | 60.58 | 3288 | 55.81 | 2923 | 49.62 | 96 | 1.63 |
| **Human-specific short tandem repeats (STR) expansions** | 4910 | 272 | 5.54 | 260 | 5.30 | 236 | 4.81 | 237 | 4.83 | 298 | 6.07 | 212 | 4.32 | 131 | 2.67 |
| **Human-specific short tandem repeats (STR) contractions** | 1464 | 1165 | 79.58 | 1115 | 76.16 | 1165 | 79.58 | 1031 | 70.42 | 927 | 63.32 | 867 | 59.22 | 85 | 5.81 |
| **Human-specific mutations associated with human-specific gene expression changes in excitatory neurons\*\*** | 517 | 263 | 50.87 | 206 | 39.85 | 347 | 67.12 | 21 | 4.06 | 25 | 4.84 | 22 | 4.26 | 0 | 0.00 |
| **Human-specific mutations associated with human-specific gene expression changes in radial glia\*\*** | 947 | 476 | 50.26 | 450 | 47.52 | 683 | 72.12 | 49 | 5.17 | 41 | 4.33 | 48 | 5.07 | 0 | 0.00 |
| **Duplicated regions in GRCh38 space defined by WSSD\*\*\*** | 7897 | 5579 | 70.65 | 4969 | 62.92 | 5692 | 72.08 | 3895 | 49.32 | 3548 | 44.93 | 3236 | 40.98 | 597 | 7.56 |

Legend: \*Results of the analyses of 10Kb regions centered at the exact sites of fixed human-specific insertions are reported; \*\*Associated with changes in gene expression in human versus chimpanzee cerebral organoids; \*\*\*WSSD, whole-genome shotgun sequence detection;

Consistent with a model of the significant contribution of the ECA’s inheritance, a majority (66%-88%) of 19,221 genomic loci harboring various classes of human-specific mutations appears highly conserved in genomes of Great Apes and Rhesus (Table 1). In contrast, only 13% of 4,910 human-specific short tandem repeats (STR) expansions’ regions are conserved. Consistent with the predominantly primate’s origins of regulatory regions harboring human-specific mutations, less than 5% of analyzed sequences appear highly conserved in the mouse genome (Table 1). Two classes of HSRS were mapped to large fractions of DNA sequences highly-conserved in genomes of various species of NHP: DNA loci harboring 49.6 to 80.6% of fixed human-specific deletions and 59.2 to 79.6% of human-specific short tandem repeats (STR) contractions were identified as highly-conserved genomic regions in genomes of different species of non-human Great Apes and Rhesus (Table 1).

Genomes of the ancestral evolutionary branch leading to human and African great apes show the significant increase in duplication activity (Marques-Bonet et al., 2009; Sudmant et al., 2013) and human-specific segmental duplications have been identified among the most promising candidate genetic loci contributing to the evolution of human-specific phenotypes (Fortna et al., 2004; Dennis et al., 2012; Charrier et al., 2012; Florio et al., 2015; Dennis et al., 2017; Kronenberg et al., 2018). It was of interest to analyze evolutionary origins of 7,987 duplicated genomic regions that were mapped to the most recent hg38 release of the human reference genome using whole-genome shotgun sequence detection (WSSD) algorithm (Kronenberg et al., 2018). Sequence conservation analyses of genomic loci duplicated in the human genome revealed that a vast majority of these regions (6,826 loci; 86.4%) are highly conserved in genomes of six non-human primates (Table 1; Supplemental Table S4; Figure 1). Interestingly, the largest fractions of both all regions successfully remapped from/to hg38 human reference genome (Table 1; Figure 1) and species-specific highly conserved regions (Supplemental Table S4; Figure 1) were observed in Gorilla and Chimpanzee genomes, indicating that these two species of non-human Great Apes are the similarly close to Modern Humans based on conservation patterns of genomic regions duplicated in the human genome. Present observations clearly demonstrate the species-specific mosaicism of evolutionary origins of genomic regions duplicated in the genome of Modern Humans (Table 1; Figure 1; Supplemental Table S4).

**Figure 1.**
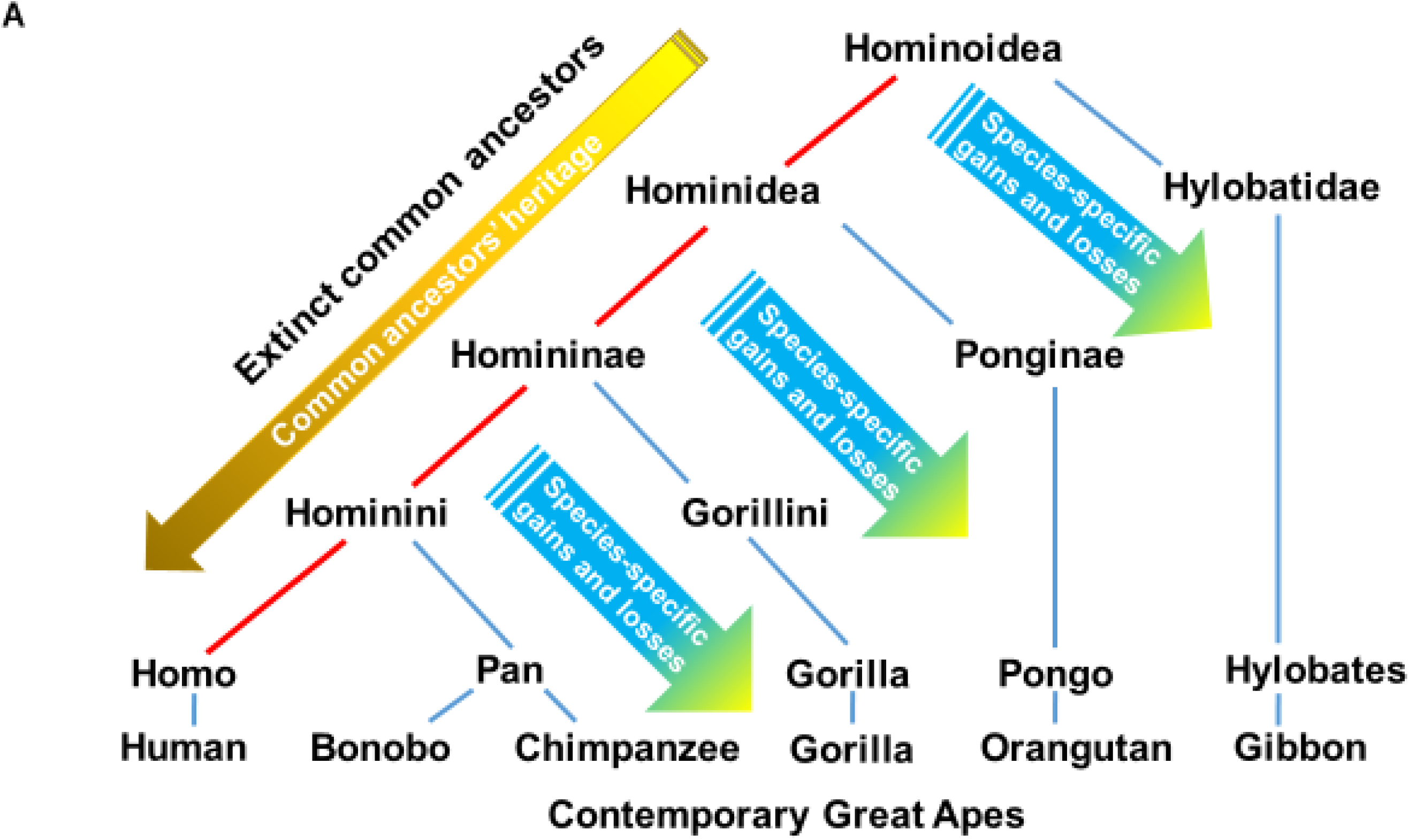
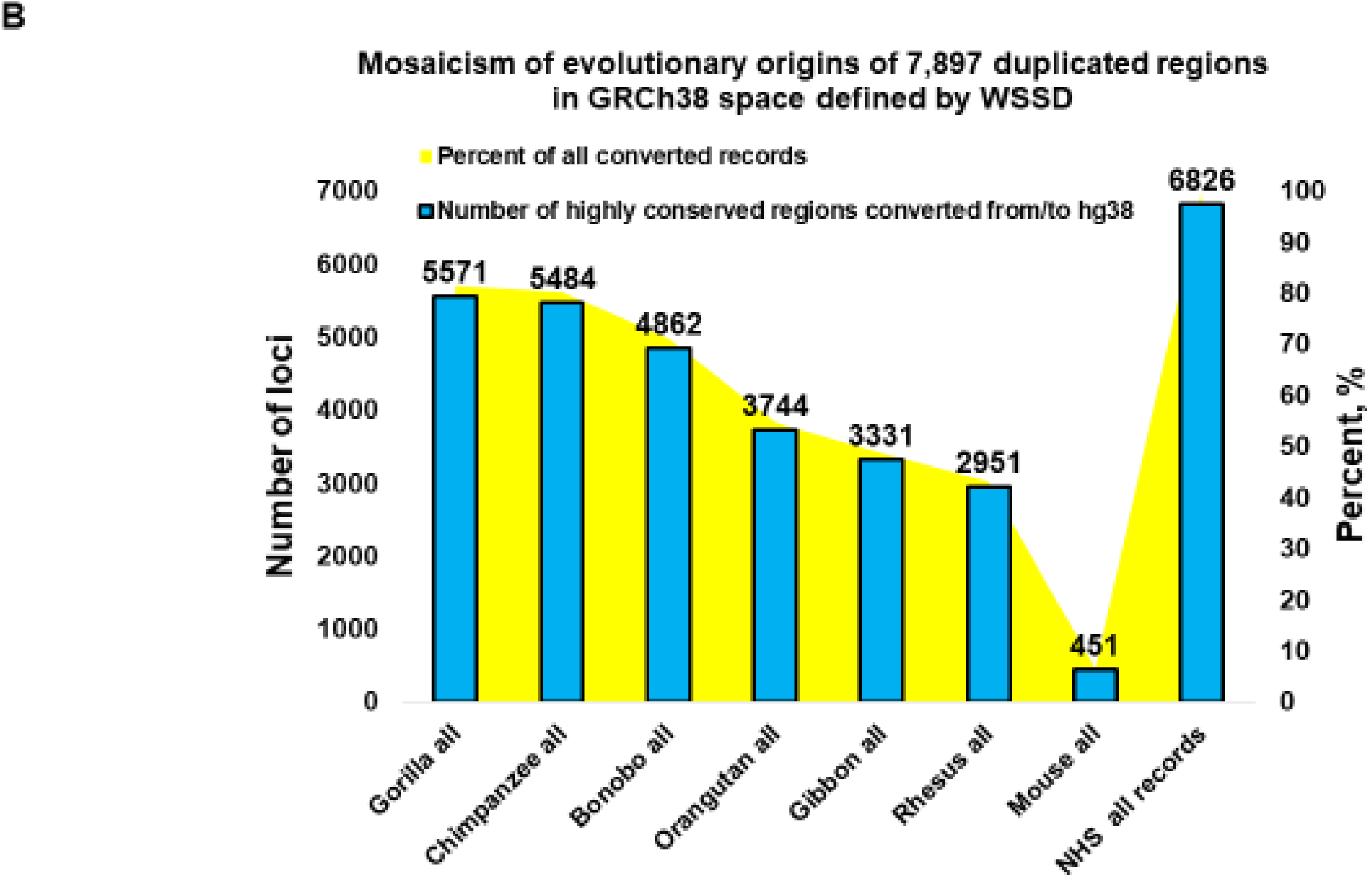
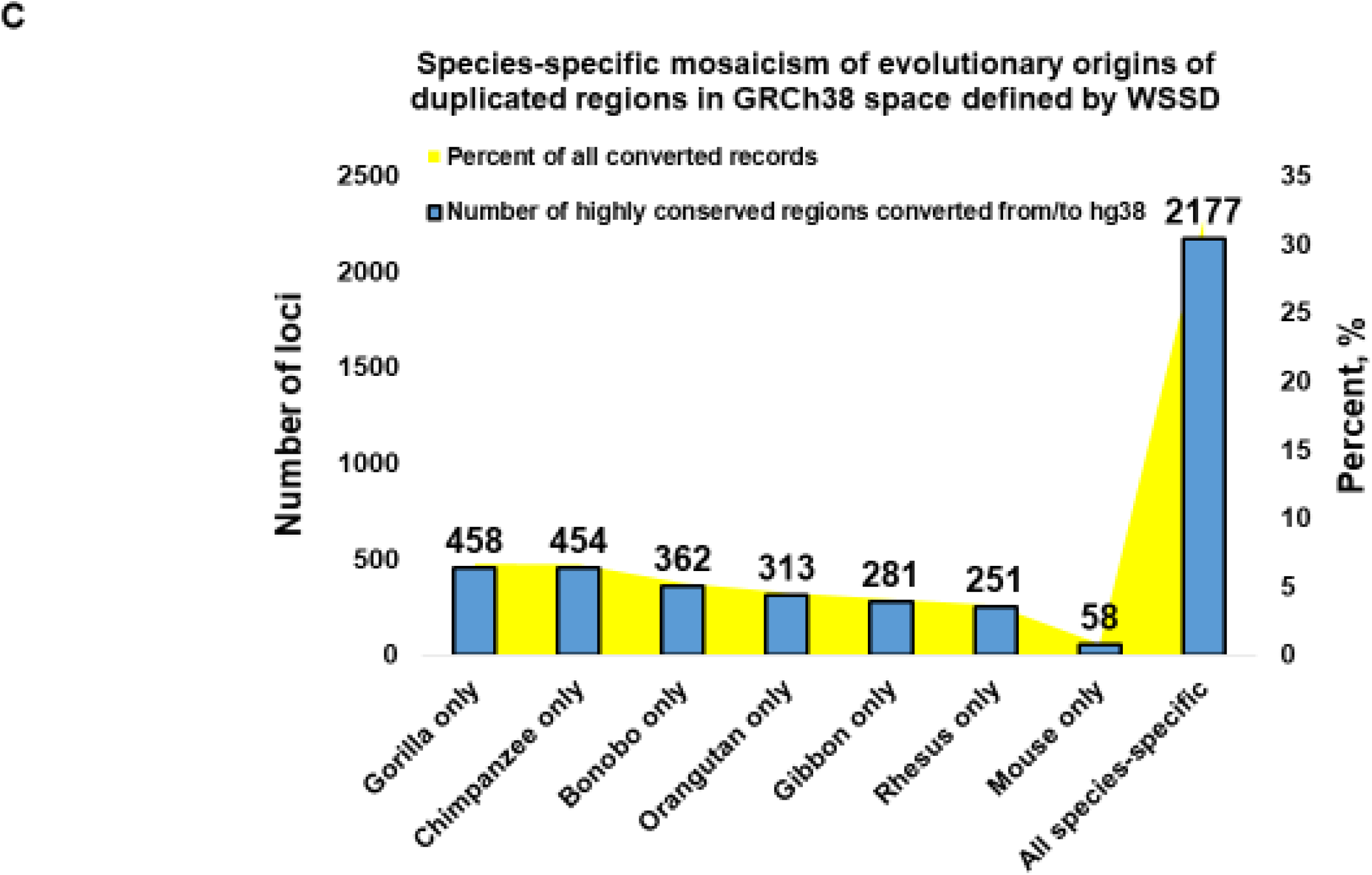
Mosaicism of evolutionary origins of 7,897 duplicated regions in the hg38 release of human reference genome defined by whole-genome shotgun sequence detection (WSSD). A. A consensus model of the lineage speciation during the evolution of Great Apes. The arrows depict the hypothetical flow of genomic information inherited from extinct common ancestors (ECAs) and acquired through species-specific gain and losses. Evolutionary origins via ECA’s inheritance pathways of highly-conserved sequences in genomes of Modern Humans and non-human species of Great Apes are postulated. B. Mosaicism of evolutionary origins of 7,897 duplicated regions in the hg38 release of human reference genome defined by WSSD (Kronenberg et al., 2018). Numbers of highly-conserved regions that successfully completed direct and reciprocal conversion tests are reported for each non-human species (NHS). C. Species-specific mosaicism of evolutionary origins of duplicated regions in the hg38 release of human reference genome defined by WSSD. Only highly-conserved sequences unique for each genome of NHS are reported.

### Mosaicism of evolutionary origins of genomic loci harboring human-specific mutations associated with human-specific gene expression changes in excitatory neurons and radial glia

Overall, the fractions of highly-conserved sequences harboring human-specific mutations and assigned to different NHP’s species seem to reflect the consensus evolutionary order of the NHP genomes’ similarity to the genome of Modern Humans. Most notable exceptions from this pattern were identified during the analyses of human-specific mutations associated with human-specific gene expression changes in excitatory neurons and radial glia (Tables 1 & 2). Among NHP species, a markedly prominent majority of genomic regions harboring human-specific mutations associated with human-specific gene expression changes during brain development in both excitatory neurons and radial glia is highly conserved only in genomes of our three closest evolutionary relatives: 50.9% and 50.3% in Chimpanzee; 39.9% and 47.5% in Bonobo; 67.1% and 72.1% in Gorilla (for excitatory neurons and radial glia, respectively). In contrast, only ∼ 5% of genomic regions harboring human-specific mutations associated with human-specific gene expression changes detected in cerebral organoids are highly-conserved in genomes of Orangutan, Gibbon, and Rhesus. Among non-human Great Apes, most significant fractions of candidate HSRS associated with human-specific gene expression changes in both excitatory neurons (347 loci; 67%) and radial glia (683 loci; 72%) are highly conserved in the Gorilla genome. For highly-conserved regions harboring HSRS associated with human-specific gene expression changes in excitatory neurons, differences in conservation profiles between genomes of Gorilla, Chimpanzee, and Bonobo were highly significant as defined by the two-tailed Fisher’s exact test (p = 1.434E-07; p = 1.574E-18; p = 0.000463). For highly-conserved regions harboring HSRS associated with human-specific gene expression changes in radial glia, differences in conservation profiles between genomes of Gorilla and Chimpanzee as well as Gorilla and Bonobo were highly significant (p = 1.623E-22 and p = 7.369E-28, respectively). In contrast, conservation profiles of HSRS associated with human-specific gene expression changes in radial glia were similar in genomes of Chimpanzee and Bonobo (p = 0.250; Table 1). These observations suggest that the majority of highly-conserved genomic regions harboring candidate HSRS associated with human-specific differences of gene expression in both excitatory neurons and radial glia was inherited from ECAs of Modern Humans and Gorilla. This conclusion remains valid when the analyses were performed considering either only loci remapped to/from NHP’s genomes to identical hg38 genomic coordinates (Table 2; Figures 2-3; Supplemental Table S4) or only genomic loci uniquely mapped to genomes of only single species of non-human Great Apes (Figure 3). Notably, differences in conservation profiles of genomic loci harboring HSRS associated with human-specific gene expression changes in radial glia appear particularly prominent (Table 2; Figures 2-3; Supplemental Table S5). Consistent with the ECA’s inheritance model, from 85.3% to 95.1% of HSRS-harboring regions that are highly-conserved in the genomes of Chimpanzee and Bonobo are highly-conserved in the Gorilla genome as well. In contrast, only from 59.0% to 59.3% and from 62.7% to 65.1% of HSRS-harboring regions that are highly-conserved in the Gorilla genome remain highly-conserved in the genomes of Bonobo and Chimpanzee, respectively. These findings were corroborated by observations demonstrating common evolutionary patterns of 248 insertions sites of African ape-specific retrovirus PtERV1 (45.9%; p = 1.03E-44) intersecting genomic regions harboring 442 HSRS, which are enriched for HSRS that have been associated with human-specific (HS) gene expression changes in cerebral organoid models (Supplemental Text; Supplemental Tables S6a; S6b; and S6c).

**Table 2.**
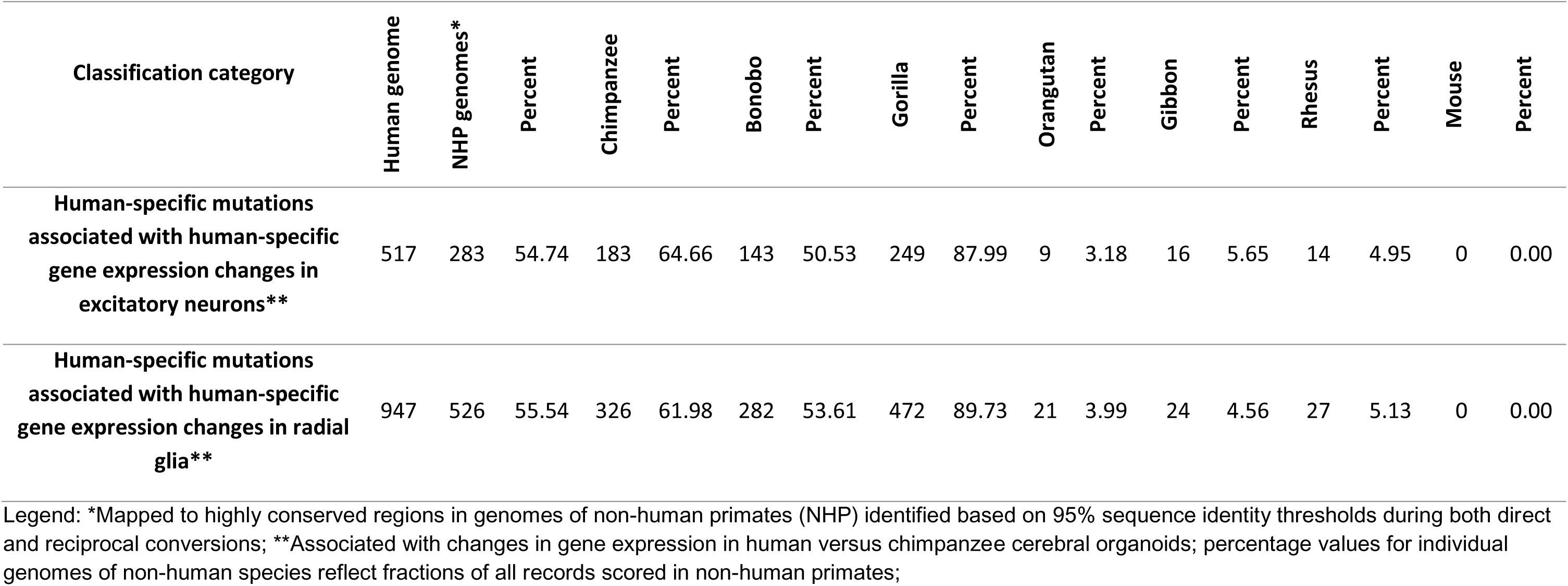
Mosaicism of evolutionary origins of genomic loci harboring candidate human-specific regulatory sequences associated with human-specific gene expression changes in human versus chimpanzee brain organoids. Only records remapped to the identical hg38 loci are reported.

**Figure 2.**
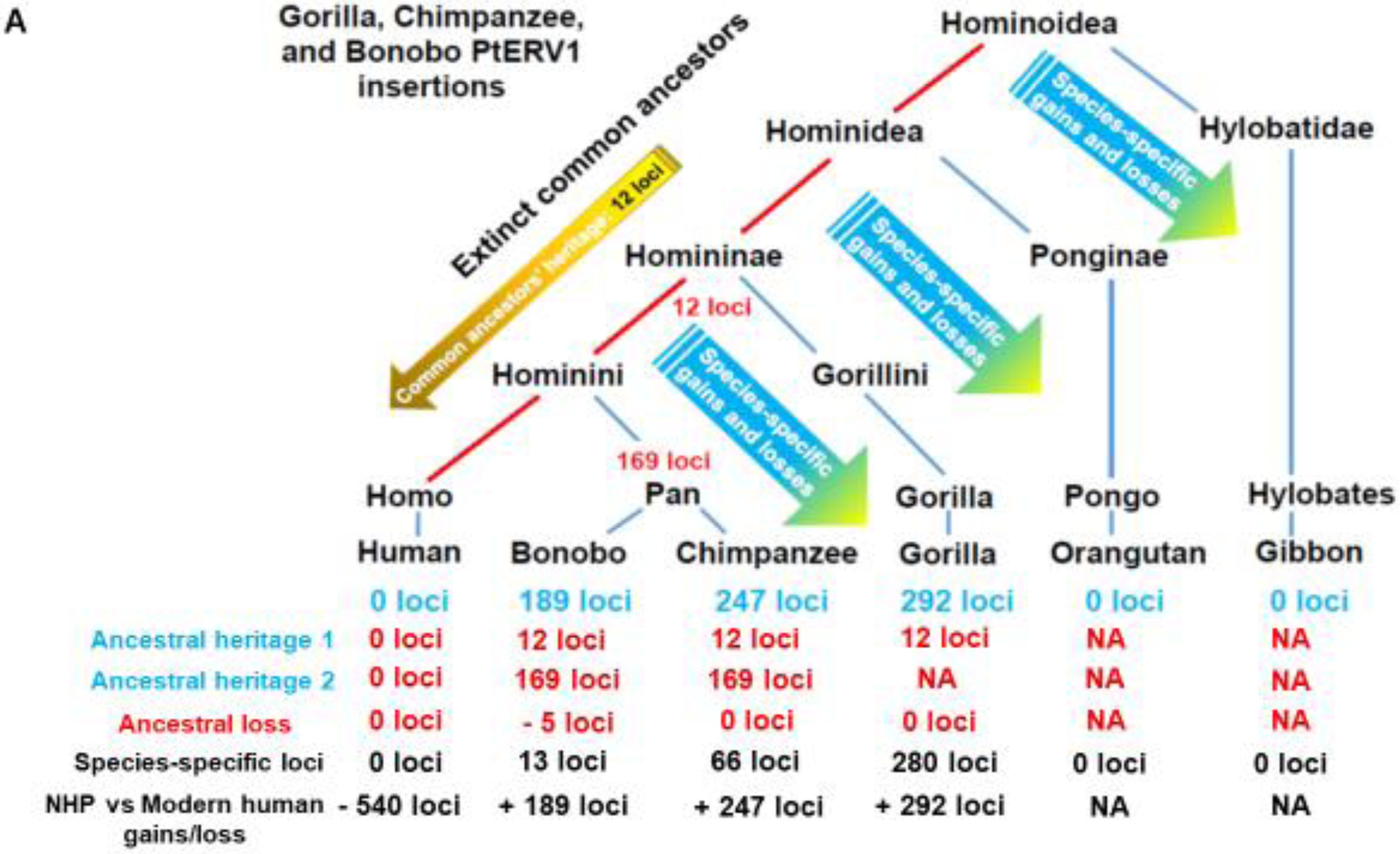
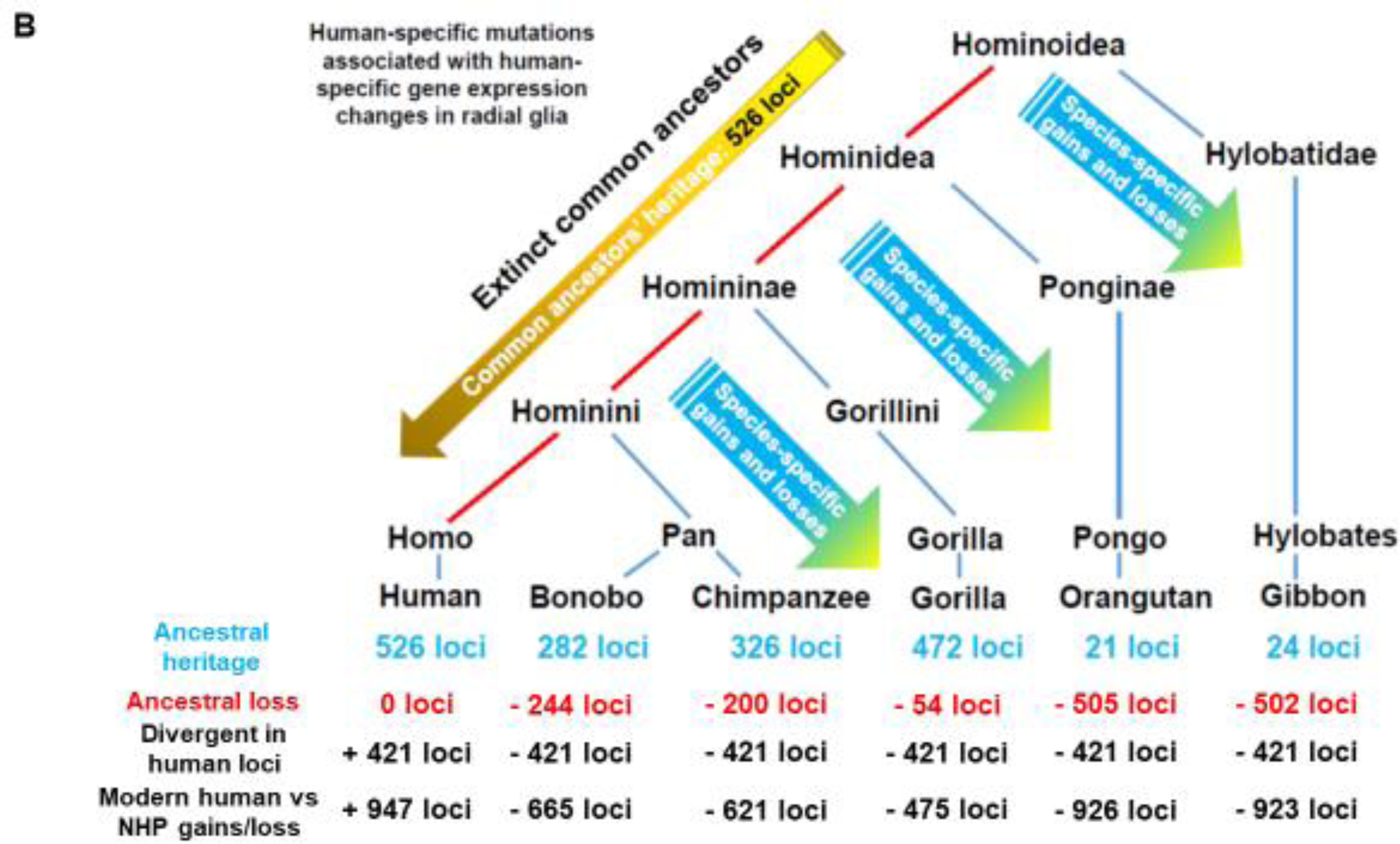
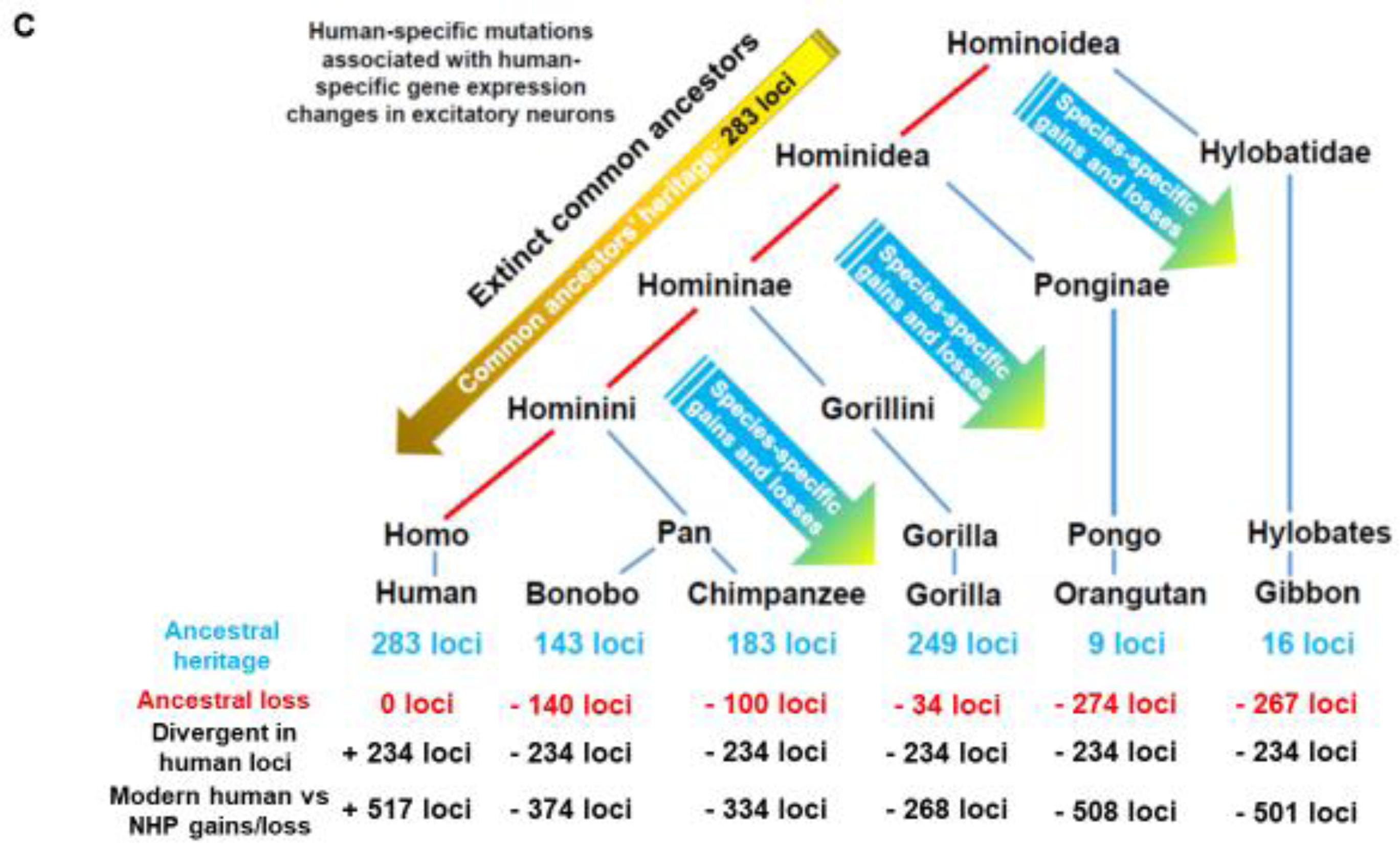
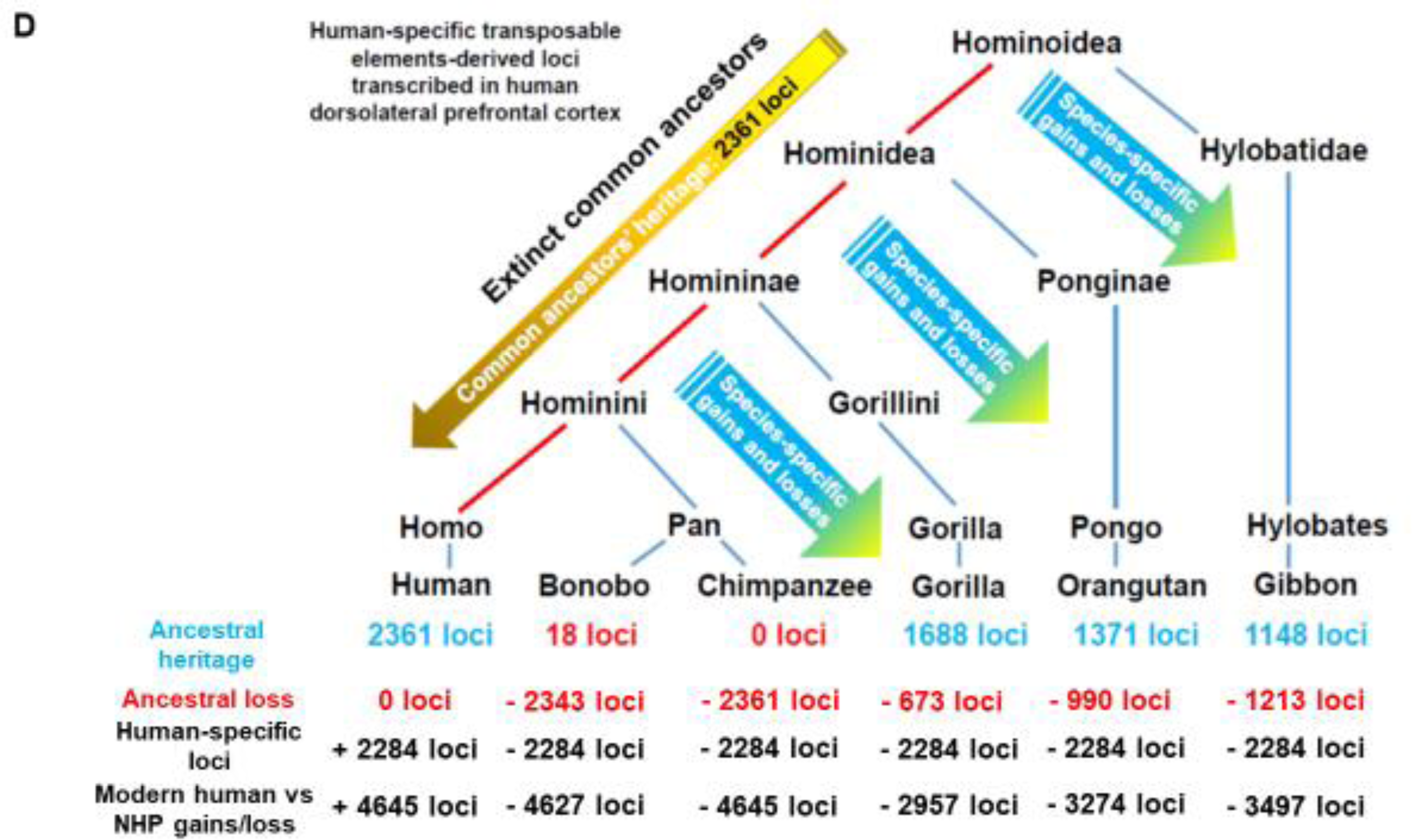
Mosaicism of evolutionary origins of candidate human-specific and primate-specific regulatory loci defined by sequence conservation analyses. A. Evolutionary patterns of inheritance, gains, and losses of 540 insertions of Africa Great Apes-specific retrovirus PtERV1. B. Evolutionary patterns of inheritance, gains, and losses of 947 human-specific regulatory regions associated with human-specific changes of gene expression in radial glia. Only records identically remapped loci in NHP during direct and reciprocal conversions from/to hg38 are reported. C. Evolutionary patterns of inheritance, gains, and losses of 517 human-specific regulatory regions associated with human-specific changes of gene expression in excitatory neurons. Only records identically remapped loci in NHP during direct and reciprocal conversions from/to hg38 are reported. D. Evolutionary patterns of inheritance, gains, and losses of 4645 human-specific transposable elements (TE) loci transcriptionally active in human dorsolateral prefrontal cortex (DLPFC). Common ancestor heritage number of 2361 loci takes into account 1,045 loci highly conserved in Rhesus; 4645 loci failed conversion to PanTro5 & PanPan1 (sequence identity threshold of 10%); 18 loci completed direct & reciprocal conversions (sequence identity threshold of 95%) from/to hg38 & PanPan2; 4612 failed conversions to PanTro5; PanPna1; PanPan2 (sequence identity threshold of 10%).

**Figure 3.**
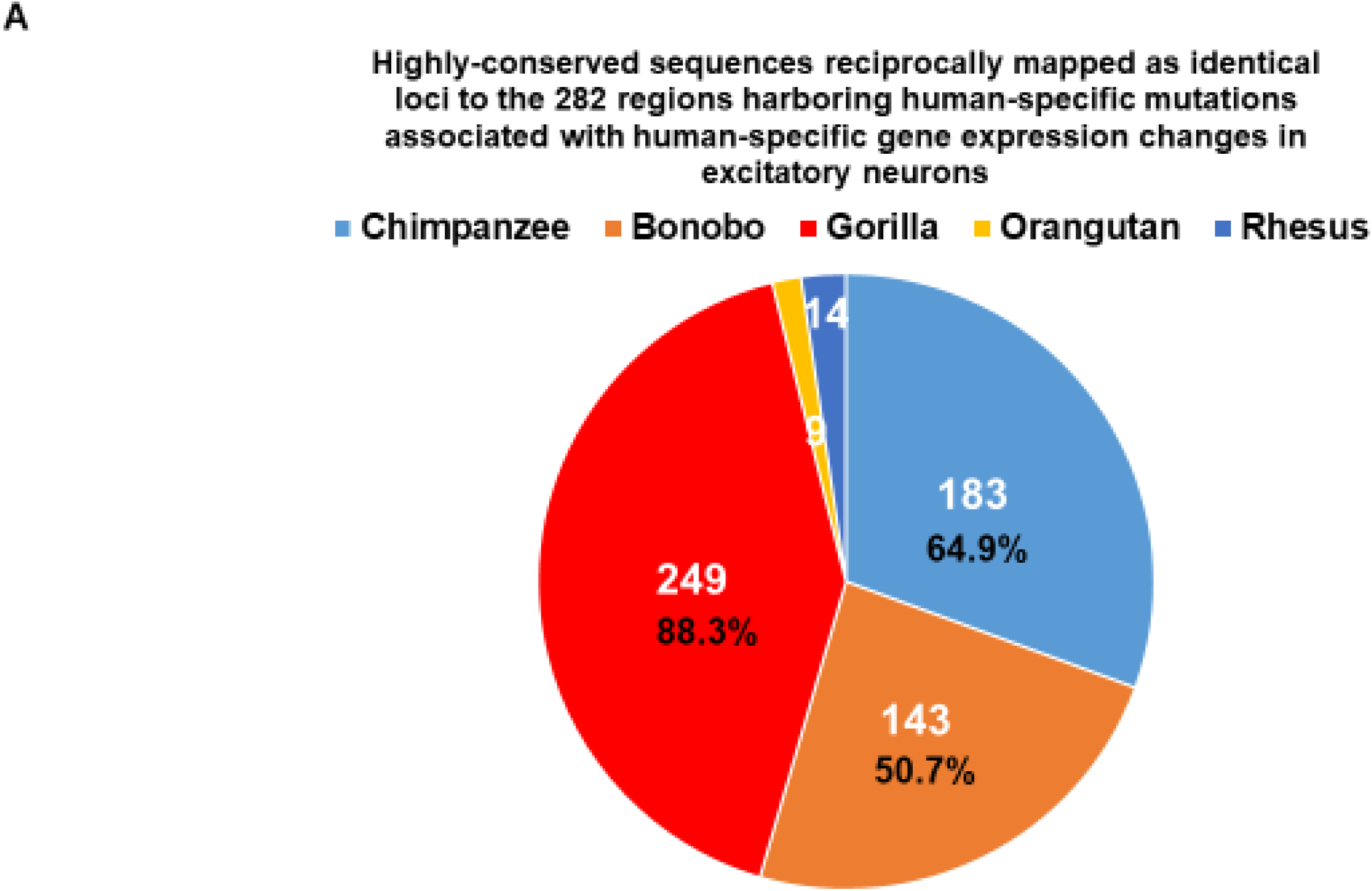
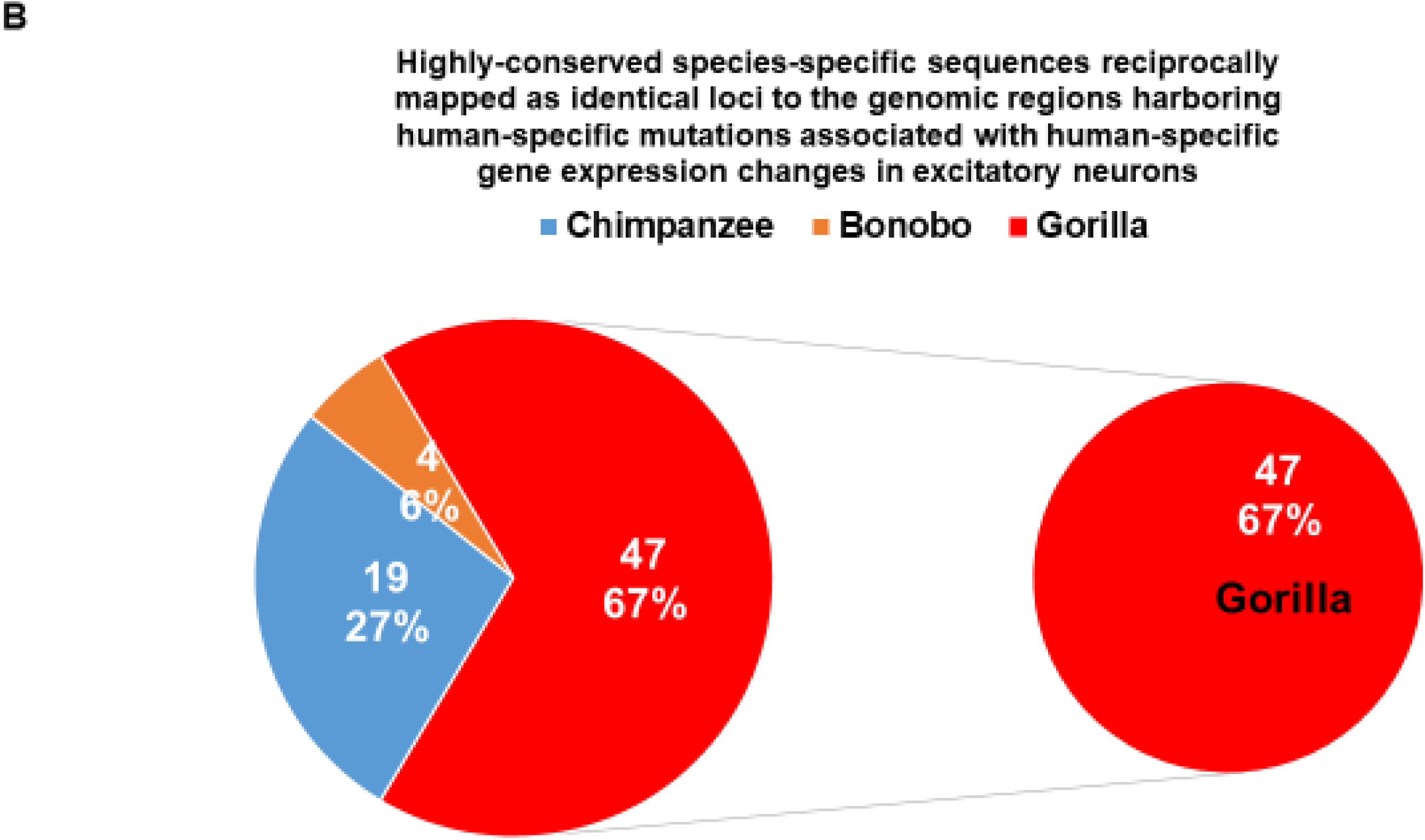
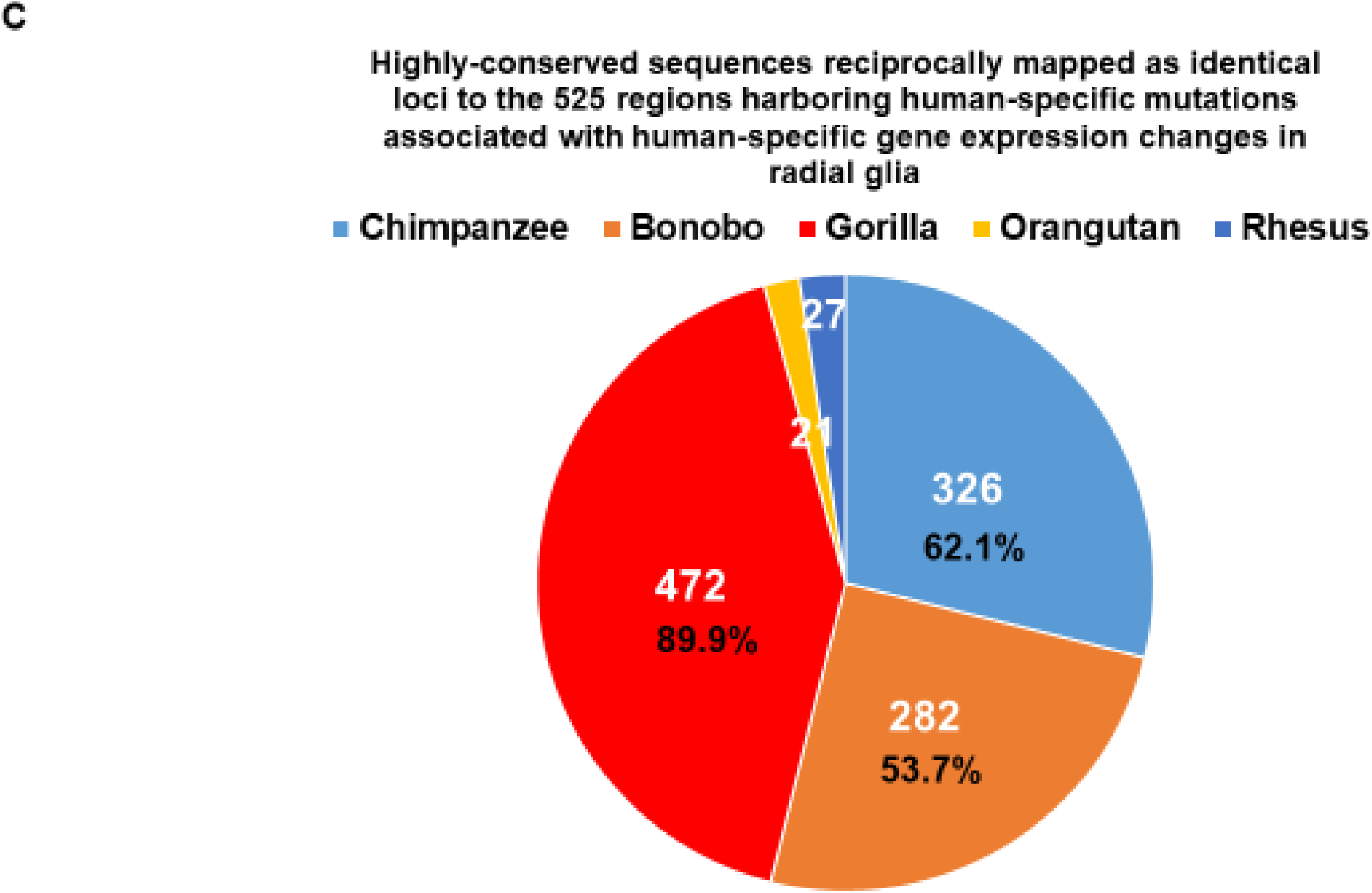
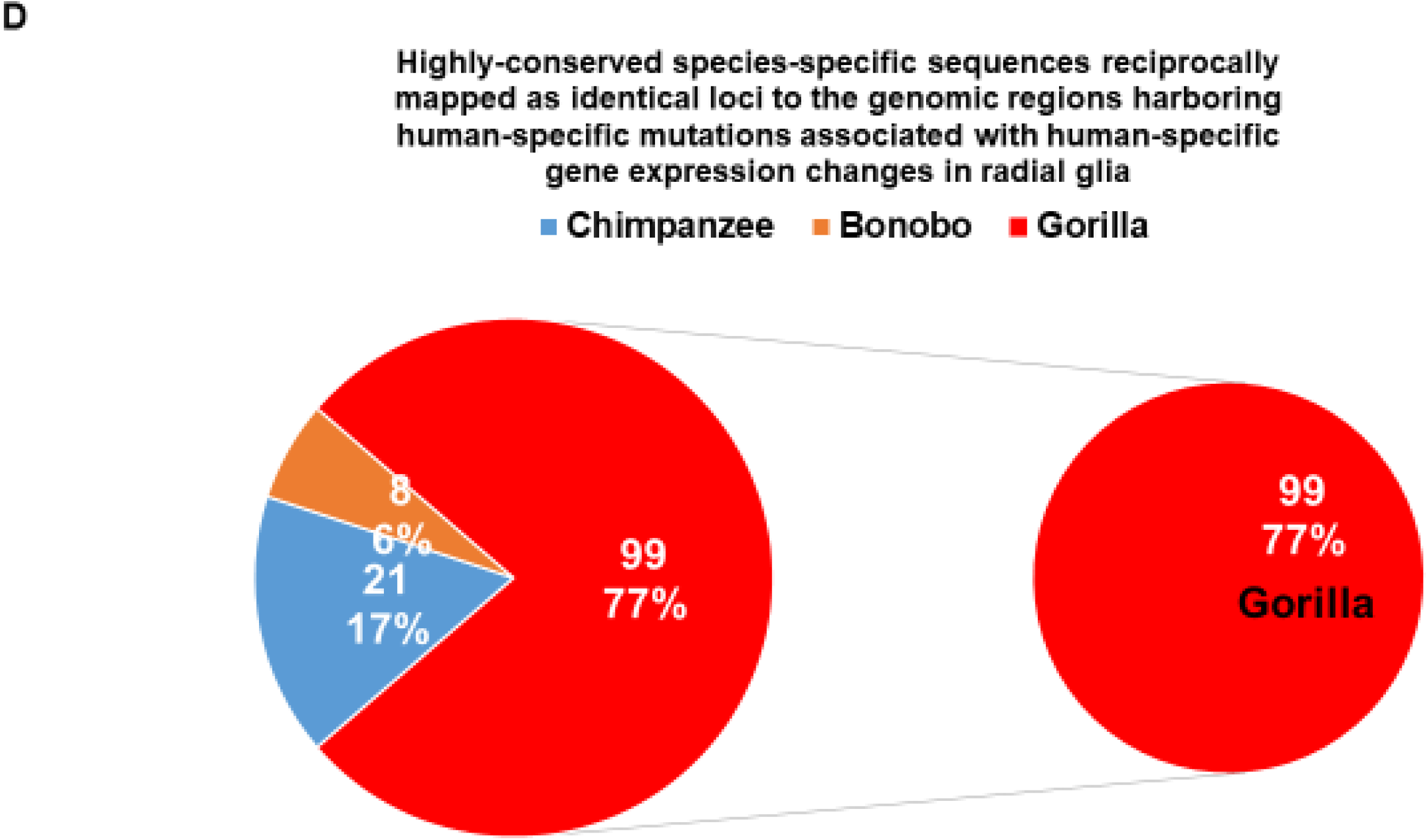
Species-specific mosaicism of evolutionary origins of human-specific regulatory sequences associated with human-specific changes of gene expression in excitatory neurons (A, B) and radial glia (C, D). A. Highly-conserved sequences reciprocally mapped as identical loci to the 282 regions harboring human-specific mutations associated with human-specific gene expression changes in excitatory neurons. B. Highly-conserved species-specific sequences reciprocally mapped as identical loci to the genomic regions harboring human-specific mutations associated with human-specific gene expression changes in excitatory neurons. C. Highly-conserved sequences reciprocally mapped as identical loci to the 525 regions harboring human-specific mutations associated with human-specific gene expression changes in radial glia. D. Highly-conserved species-specific sequences reciprocally mapped as identical loci to the genomic regions harboring human-specific mutations associated with human-specific gene expression changes in radial glia.

### Mosaicism of evolutionary origins of candidate human-specific regulatory loci defined based on the mapping failure to both Chimpanzee and Bonobo reference genomes

One of the approaches to the identification of candidate HSRS is based on their absence in the genomes of our closest evolutionary relatives, Chimpanzee and Bonobo. Present analyses demonstrate that large fractions of genomic regions harboring candidate HSRS are highly-conserved in genomes of our more distant evolutionary relatives, Gorilla, Orangutan, Gibbon, and Rhesus (Tables 1-2). These observations suggest that candidate human-specific regulatory loci which were defined based on the mapping failures to both Chimpanzee and Bonobo genomes may originate on DNA sequences highly-conserved in genomes of other NHP species. To test the validity of this hypothesis, 16,730 genomic loci harboring distinct classes of HSRS were identified that failed to convert to genomes of both Chimpanzee and Bonobo using 10% sequence identity threshold (Table 5). Then highly-conserved sequences in genomes of Rhesus, Gibbon, Orangutan, and Gorilla were identified and tabulated for each category of HSRS. Consistent with the bypassing patterns of evolutionary inheritance, thousands of distinct classes of candidate HSRS that failed to map to genomes of both Chimpanzee and Bonobo are highly conserved in genomes of Gorilla, Orangutan, Gibbon, and Rhesus (Table 5). Most prominently, significant fractions of retrotransposon-derived loci that are transcriptionally-active in human dorsolateral prefrontal cortex and absent in genomes of both Chimpanzee and Bonobo are highly conserved in genomes of Gorilla, Orangutan, Gibbon, and Rhesus (1,688; 1,371; 1,148; and 1,045 loci, respectively; Figure 4). It has been observed that in all instances the Gorilla genome had the largest numbers of shared with Modern Humans highly-conserved sequences that failed to map to genomes of both Chimpanzee and Bonobo (Supplemental Table S7).

**Figure 4.**
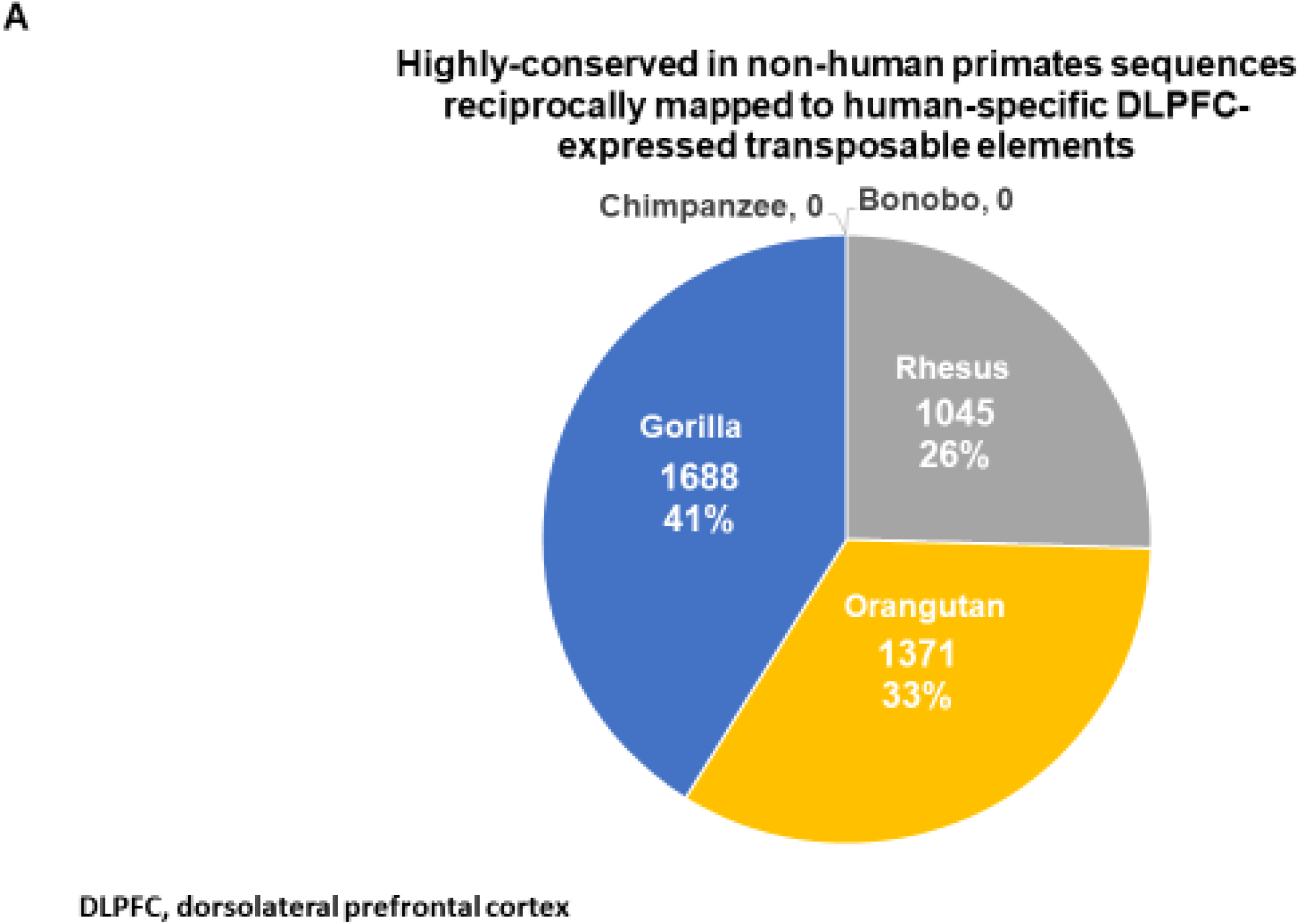
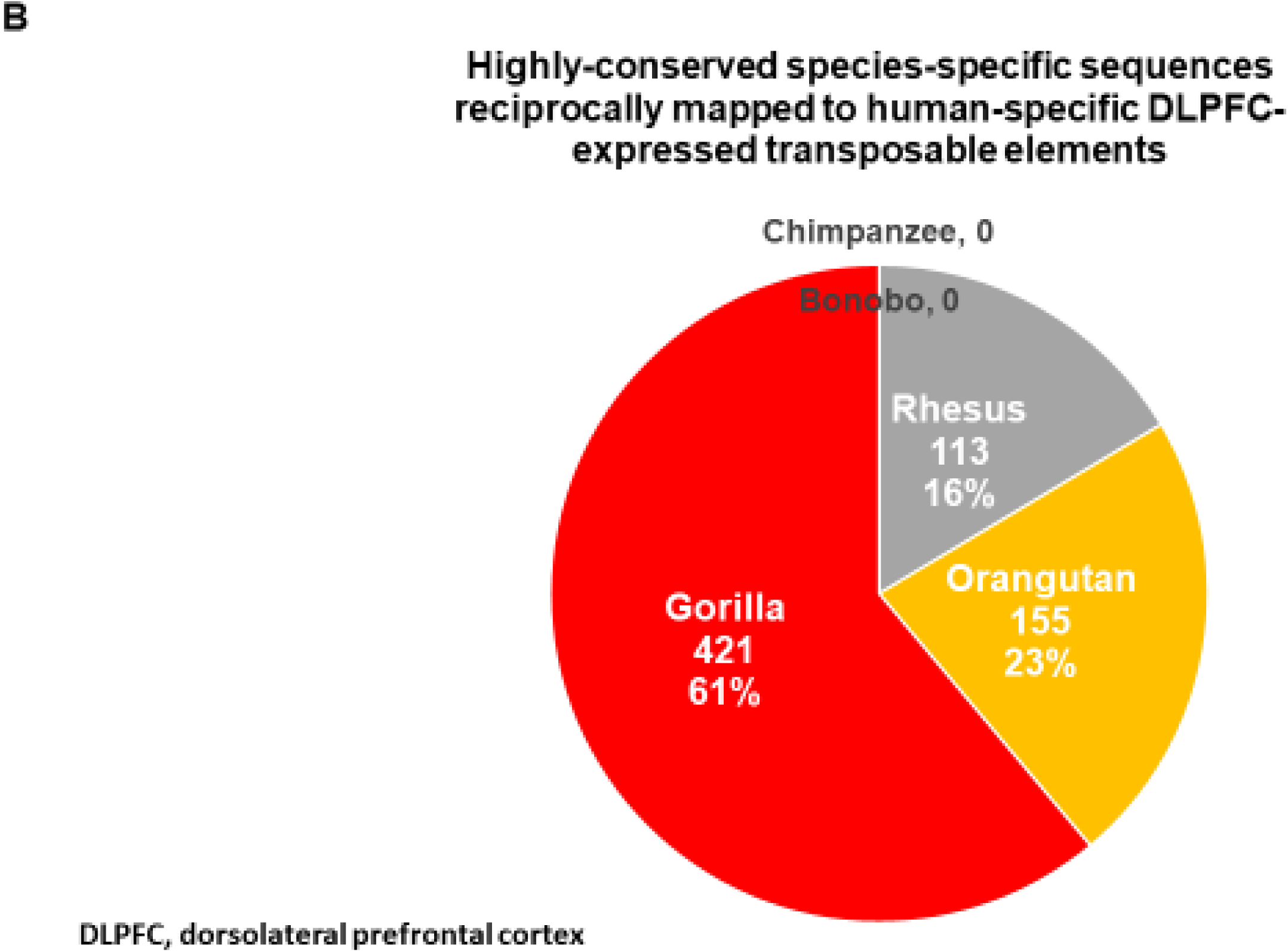
Species-specific mosaicism of evolutionary origins of human-specific regulatory sequences encoded by transcriptionally-active transposable elements (TE) in human DPLFC. A. Numbers of sequences highly-conserved in NHP genomes reciprocally mapped to human-specific DLPFC-expressed transposable elements. B. Highly-conserved species-specific sequences reciprocally mapped from NHP genomes to human-specific DLPFC-expressed transposable elements.

These observations indicate that a more stringent approach for definition of candidate HSRS which are likely to have been created de novo in the genome of Modern Humans would be to require the conversion failures of a regulatory DNA sequence to all six NHP’s genomes, namely genomes of Chimpanzee, Bonobo, Gorilla, Orangutan, Gibbon, and Rhesus. Using this strategy, 12,486 candidate HSRS have been identified (Supplemental Table S8), indicating that 24.8% of all analyzed in this contribution HSRS-harboring genomic loci could be classified as created de novo candidate HSRS.

### Enrichment within human-specific regulatory pathways of genes comprising expression signatures of human-specific neurodevelopmental and pluripotency transcriptional networks

It is possible that distinct types of candidate HSRS are not just unrelated elements of a random population of DNA sequences, but they might represent a coherent collection of regulatory DNA sequences assembled during evolution to facilitate execution of human-specific functions. If this hypothesis is correct, then HSRS may represent the key components of human-specific genomic regulatory pathways governing human-specific gene expression patterns observed in various phenotypic contexts associated with human-specific traits. Therefore, the validity of this hypothesis could be tested using strictly-defined by comparisons with non-human Great Apes human-specific gene expression signatures associated with development of human brain in the cerebral organoid model (Kronenberg et al., 2018) and induced pluripotency phenotypes of human versus NHP cells (Marchetto et al., 2013). Kronenberg et al. (2018) identified several hundred genes that manifest human-specific expression changes in Modern Humans versus Chimpanzee cerebral organoids’ model of brain development. Significant fractions of these human-specific neurodevelopmental networks operating in both excitatory neurons and radial glia appear associated with human-specific structural variants, specifically, with human-specific insertions and deletions (Supplemental Table S9). Even larger fractions of genes comprising human-specific neurodevelopmental networks have been identified as components of the gene expression signature (GES) of the MLME cells in human preimplantation embryo: common gene sets represent 262 genes (68.2%; p = 1.59E-93) and 481 genes (72.0%; p = 3.95E-187) for human-specific GES of excitatory neurons and radial glia, respectively (Supplemental Table S9).

It has been reported that the creation of the MLME cells in human preimplantation embryos is associated with increased expression of primate-specific retrotransposon-derived regulatory long non-coding RNAs termed human pluripotency-associated transcripts, HPATs (Glinsky et al., 2018). Most recently, the expansive networks of primate-specific and human-specific retrotransposons transcriptionally active in human dorsolateral prefrontal cortex (DLPFC) and associated coding genes have been identified (Guffanti et al., 2018). Thus, it was of interest to determine what fractions of genes comprising human-specific neurodevelopmental gene expression networks in excitatory neurons and radial glia may overlap with coding genes coupled with active transcription of transposable elements in human DLPFC. Remarkably, it has been observed that common gene sets represent a vast majority of genes comprising human-specific neurodevelopmental networks: they comprise 322 genes (83.9%; p = 5.57E-84) and 561 genes (84%; p = 3.08E-146) for human-specific GES of excitatory neurons and radial glia, respectively (Supplemental Table S9). Interestingly, both *SRGAP2C* and *ARHGAP11B* genes driving divergent cortical development between humans and chimpanzee (Dennis et al., 2012; Charrier et al., 2012; Florio et al., 2015) harbor transposable elements transcriptionally active in human DLPFC (Guffanti et al., 2018).

Marchetto et al. (2013) identified human-specific gene expression signature distinguishing induced pluripotent stem cells (iPSC) engineered from cells of Modern Humans and NHP species. To infer the putative regulatory association patterns of genes distinguishing human iPSC versus NHP iPSC and human-specific genomic regulatory pathways, overlap enrichment analyses of corresponding gene sets have been performed. It has been observed, that the association patterns of human-specific gene expression signatures of brain development and pluripotency phenotypes with compendiums of genes likely governed by human-specific regulatory pathways appear strikingly similar (Supplemental Tables S9 and S10). Overall, 88% of genes comprising human-specific expression signature of the induced pluripotency phenotype represent genes implicated in putative regulatory associations with human-specific genomic pathways. These observations support the hypothesis that human-specific gene expression signatures of brain development and pluripotency phenotypes are associated with a collection of HSRS assembled during evolution into human-specific genomic regulatory pathways to govern transcriptional networks in human cells.

### Enrichment within human-specific regulatory pathways of genes encoding virus-interacting proteins

Present analyses identified thousands of genes that are likely regulated by HSRS, among which *SERINC5*, *APOBEC3B*, and *PIWIL2* genes are of particular interest because high expression of proteins encoded by these genes is likely to confer increased resistance to the retroviral infection and to restrict the propagation of retrotransposons in human cells (Usami et al., 2015; Rosa et al., 2015; Marchetto et al., 2013). These observations suggest that genes involved in interactions between viruses and human cells may constitute an important category of genetic targets that are placed under control of HSRS during speciation of the human lineage. Therefore, it was of interest to determine whether human genes encoding virus-interacting proteins (VIPs) are components of human-specific regulatory networks by analyzing a comprehensive set of more than four thousand genes encoding human VIPs recently reported by Enard and Petrov (2018). The set of VIP-encoding genes analyzed in this study comprised of 4,433 individual records, which were obtained after removal of three loci that are no longer in the current ENSEMBL database and adjustment for records that have multiple ENSEMBL identifying numbers associated with the same genomic loci defined by unique gene symbols in hg38 release of the human reference genome database. It has been observed that a significant fraction of genes encoding human VIPs (27.2%; p = 1.03E-66; the hypergeometric test) appear associated with human-specific structural variants, specifically, with human-specific insertions and deletions (Supplemental Table S11). A prominent majority of genes encoding human VIPs have been identified as components of the expression signature of the MLME cells in human preimplantation embryo (3,551 VIPs; 80.1%; Supplemental Table S11) and coding genes coupled with active transcription of transposable elements in human DLPFC (3,574 VIPs; 80.6%; Supplemental Table S11). Intriguingly, the *LBP9/HERVH* gene expression pathway regulated in hESC by primate-specific retrovirus LTR7/HERVH-derived long-non coding RNAs appears to control the expression of 2,661 VIPs (60%; Supplemental Table S11). Overall, a vast majority of genes encoding human VIPs (4,253 of 4,433 genes; 95.9%) has been identified as putative downstream targets of human-specific genomic regulatory networks. Importantly, these regulatory networks were defined employing vastly different experimental, analytical, and computational approaches that were applied within the broad range of experimental settings: i) Great Apes’ whole-genome sequencing-guided identification of human-specific insertions and deletions (Kronenberg et al., 2018); ii) genome-wide analysis of retrotransposon’s transcriptome in postmortem samples of human dorsolateral prefrontal cortex (Guffanti et al., 2018); iii) shRNA-mediated silencing of LTR7/HERVH retrovirus-derived long non-coding RNAs in hESC (Wang et al., 2014); iv) single-cell expression profiling analyses of human preimplantation embryos (Glinsky et al., 2018).

### VIP-encoding genes represent large fractions of genes comprising human-specific expression signatures of radial glia, excitatory neurons, and induced pluripotent stem cells

Above considerations suggest that VIP-encoding genes may be represented among genes comprising human-specific expression signatures of neurodevelopmental and pluripotency networks (see above). Follow-up analyses revealed that VIPs indeed constitute large fractions of genes comprising human-specific expression signatures of radial glia (34%; p = 1.51E-94; the hypergeometric test; Figure 5), excitatory neurons (30%; p = 1.92E-41; Figure 5); and human induced pluripotent stem cells (25%; p = 1.22E-08). Intriguingly, disproportionally large fractions of VIP-encoding genes comprising human-specific expression signatures of both up-regulated and down-regulated genes in radial glia appear associated with human-specific structural variants (Figure 5; Table 5). It has been observed that the association with human-specific mutations of VIPs is 6.5-fold greater compared to non-VIPs (p = 1.36E-15; two-tail Fisher’s exact test) for genes up-regulated in human radial glia cells (Figure 5; Table 5). For genes down-regulated in human radial glia cells (Figure 5; Table 5), the association with human-specific mutations of VIPs is 2.1-fold greater compared to non-VIPs (p = 0.002; two-tail Fisher’s exact test). In striking contrast, no increased associations with human-specific mutations have been detected for VIP-encoding genes comprising human-specific expression signatures of excitatory neurons (data not shown), suggesting that observed putatively regulatory associations might be specific to human radial glia cells. Collectively, observations reported in this contribution are highly consistent with the hypothesis that VIP-encoding genes represent important components of human-specific regulatory networks operating in phenotypically distinct human cells which are essential for biological functions of cognition and pluripotency.

**Figure 5.**
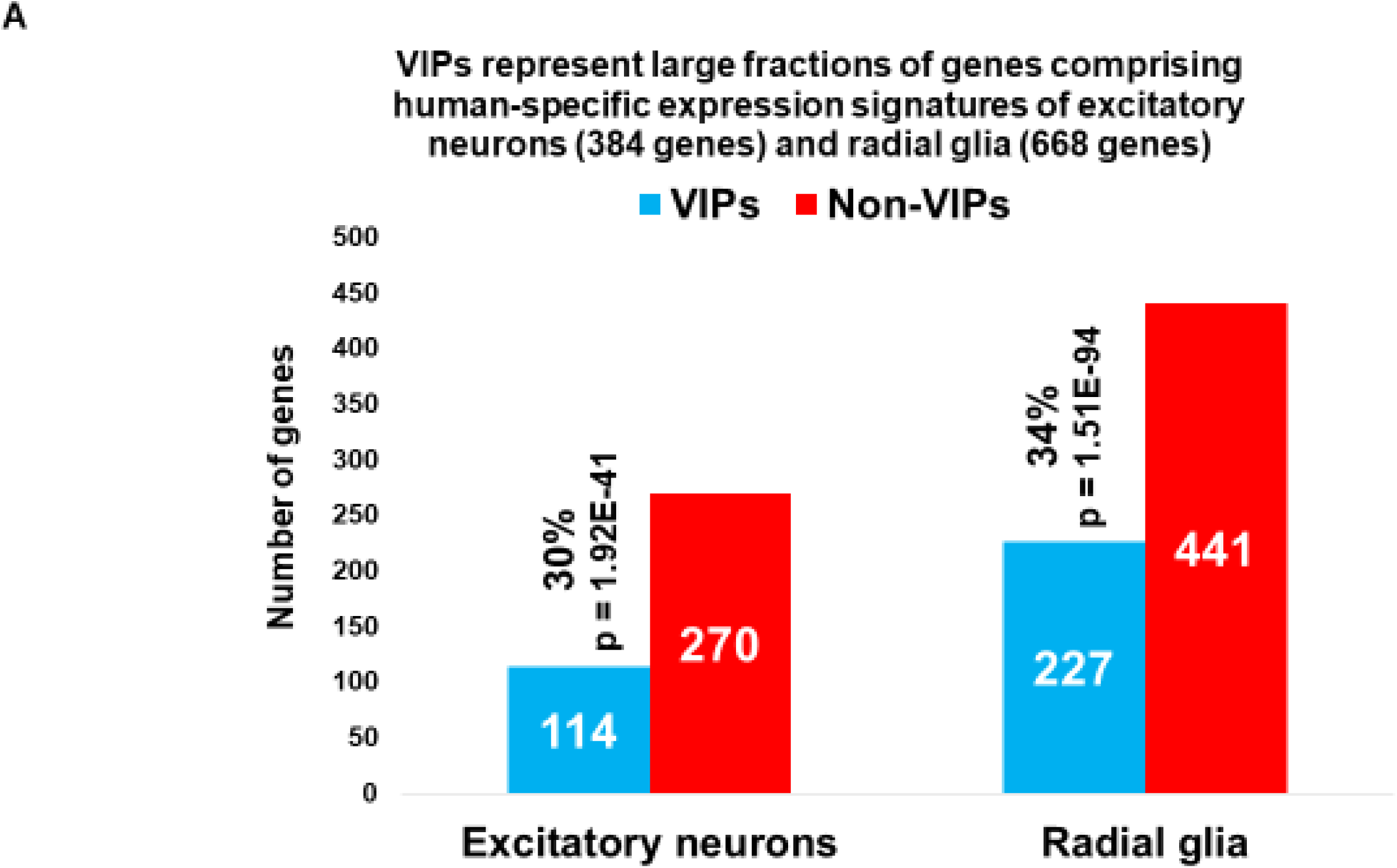
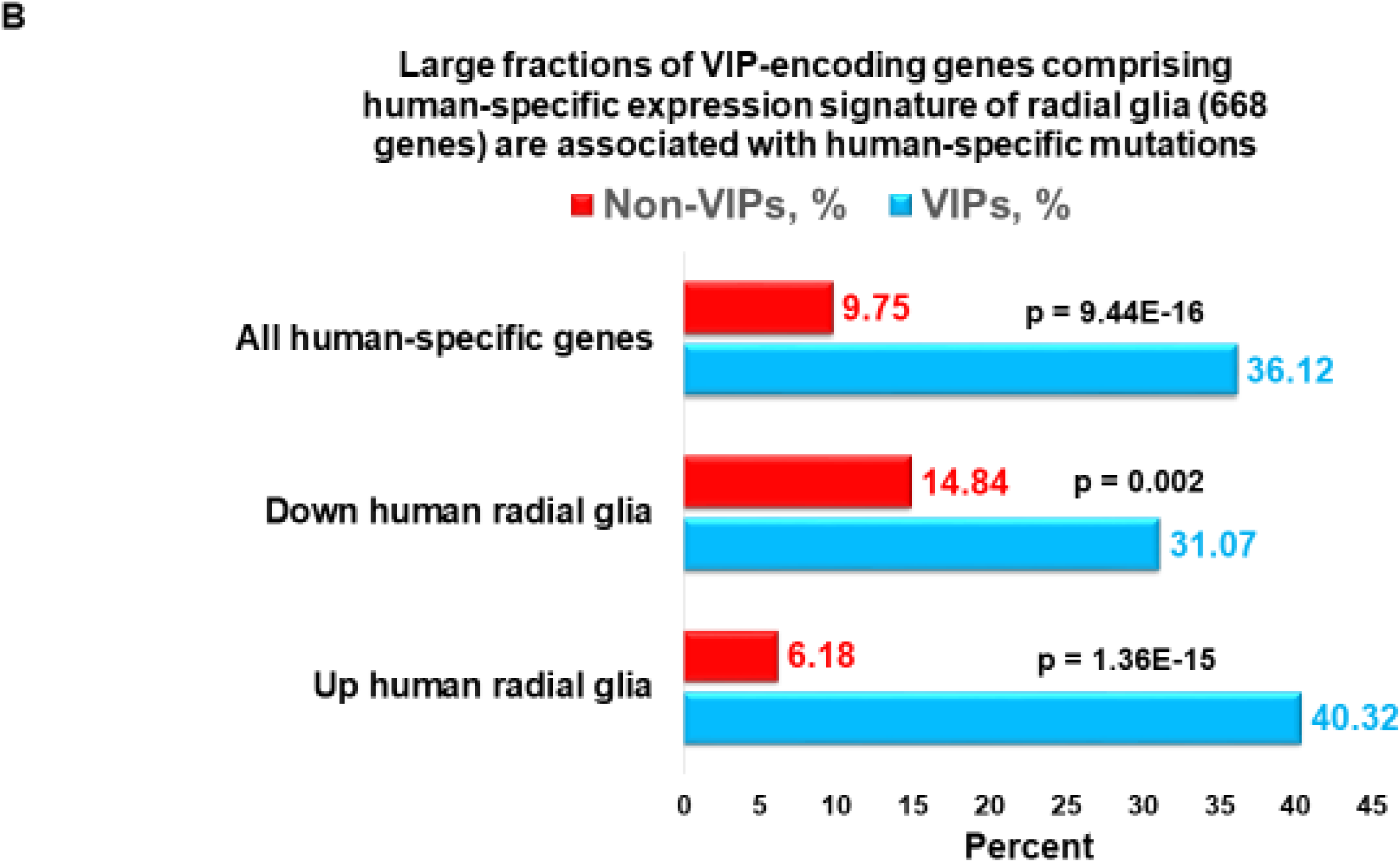
Genes encoding virus-interacting proteins (VIPs) represent large fractions of genes comprising human-specific expression signatures of excitatory neurons and radial glia. A. VIP-encoding genes comprise 34% and 30% of genes differentially-regulated in human versus chimpanzee radial glia cells and excitatory neurons, respectively, isolated from species-specific cerebral organoids. Significance of the overlaps was estimated using the hypergeometric distribution test. B. Disproportionally large fractions of both up-regulated and down-regulated VIP-encoding genes comprising human-specific expression signatures of radial glia cells are associated with human-specific mutations. Corresponding cohorts of genes encoding VIPs and non-VIPs transcripts associated with human-specific mutations were identified from the set of 668 genes manifesting human-specific expression changes in human versus chimpanzee radial glial cells (Kronenberg et al., 2018). Significance of the differences in associations with human-specific mutations between VIPs and non-VIPs genes was estimated using the two-tailed Fisher’s exact test.

## Discussion

### Implications for future structural-functional validation studies of regulatory effects of HSRS on human-specific regulatory networks and phenotypic traits

An impressive contemporary collection of nearly sixty thousand candidate HSRS assembled by the collective decades-long effort of many laboratories (see Introduction) lends further credence to the idea that unique to human phenotypes might result from human-specific changes to genomic regulatory sequences (King and Wilson, 1975). This study identifies multiple high-priority candidate HSRS for in depth structural-functional validation analyses, among which most prominent candidate HSRS appears associated with human-specific gene expression changes in excitatory neurons and radial glia as well as in human induced pluripotent stem cells. This high-priority set of elite genetic targets include candidate HSRS putatively regulating expression of *SERINC5*, *APOBEC3B*, and *PIWIL2* genes, high expression of which in human cells is likely to confer increased resistance to the retroviral infection and propagation of retrotransposons (Marchetto et al., 2013; Usami et al., 2015; Rosa et al., 2015). It is tempting to speculate that these changes may have been significant genetic contributors conferring the selective fitness advantage to human lineage during primate evolution.

Observations reported in this contribution have important implications for future functional validation studies, in particular, our approaches to selection and design of experimental systems for follow-up structural-functional analyses of the effects of specific HSRS on defined human-specific phenotypes. These considerations must take into account the history of evolutionary origins of HSRS to design the specifically tailored experimental models from a panel of cells obtained from Modern Humans and the appropriate NHP species, genomes of which either have (thus, are appropriate for targeted knock-out and silencing experiments) or not have (thus, are appropriate for knock-in and overexpression experiments) of genetic loci of interest. Results of present analyses strongly argue that these experimental systems for targeted interrogation of human-specific genomic regulatory networks should include iPSC as well as brain development models of in vitro induced iPSC differentiation and cerebral organoids. Ideally, these experimental systems should be based on cells obtained from Modern Humans, our evolutionary closest relatives, Chimpanzee and Bonobo, as well as Gorilla, Orangutan, Gibbon, and Rhesus.

### Analyses of evolutionary origins of genomic loci harboring HSRS revealed evidence of a striking ancestral polymorphism and regions of genomic divergence of Modern Humans and Great Apes

High-confidence human-specific mutations leading to emergence of candidate HSRS should be considered as rare genomic events that unlikely to occur more than once during evolution at the same genomic locations. Therefore, observations that large fractions of genomic regions harboring distinct classes of HSRS represent DNA sequences highly-conserved in genomes of distinct NHP species should be interpreted as strong circumstantial evidence consistent with their putative regulatory functions. Collectively, the evidence presented in this contribution revealed a complex unique-to-human mosaic of regulatory DNA sequences inherited from ECAs following separation events from multiple distinct NHP species and reflecting the striking ancestral polymorphism of Modern Humans. One of the novel mechanisms that may have contributed to divergence of genomic regulatory networks of Modern Humans and non-human Great Apes is illustrated by observations that the insertions of the African Great Ape-specific retrovirus PtERV1 and distinct classes of HSRS have common genomic coordinates within orthologous genomic regions of Gorilla, Chimpanzee, Bonobo, and Modern Humans. Overall, these common patterns of species-specific mutations within overlapping genomic regions were observed for nearly half (248 of 540 loci) of all PtERV1 insertions and 442 HSRS, including 21 HSRS associated with genes differentially expressed in human versus chimpanzee cerebral organoid models of brain development.

### Genomic divergence and speciation of Modern Humans reflect a complex continuing evolutionary process rather than a singular instantaneous event

Observations reported herein support the hypothesis that the speciation process during evolution of Great Apes is not likely to occur as an instantaneous event (Patterson et al., 2006): for example, human and chimpanzee lineages could have exchanged genes following the iterative sequences of the initial lineage divergence, separation, and gathering together prior to the permanent segregation of two species. Since human, chimpanzee, and gorilla lineages may have diverged during the relatively short evolutionary time, this model might reflect the extended complex speciation process of these three closely-related Great Apes, possibly involving co-evolution of their ECAs (Supplemental Figure S1). Incomplete lineage sorting events were intrinsic components of the genomic divergence and are likely played an important role in the lineage segregation. In agreement with this hypothesis, comparative analyses of multiple alignments of sequences of human, chimpanzee, gorilla, and orangutan genomes have demonstrated that a considerable fraction of genes in the human genome is more similar to the gorilla genome than to the chimpanzee genome (Patterson et al., 2006; Chen and Li, 2001; Yang, 2002; O’hUigin et al., 2002; Wall, 2003; Hobolth et al. 2007). Genomic regions harboring HSRS appear to follow the similar evolutionary trajectory. Observed examples of the bypassing pattern of the evolutionary inheritance highlight HSRS supporting the inference of alternative genealogies (human being most closely related to NHP other than Chimpanzee) are most likely reflect the incomplete lineage sorting events. Incomplete lineage sorting has been consistently observed in the multiple alignments of the genomes for human, chimpanzee, gorilla, and orangutan where differences in models of gene genealogies and species phylogeny were documented for up to 36% of the human autosomal genome (Chen and Li, 2001; Yang, 2002; Wall, 2003; Patterson et al. 2006; Hobolth et al., 2007; 2011; Kronenberg et al., 2018). Similar changes could result from species-specific losses of conserved ancestral loci of regulatory DNA, a mechanism that contributed to evolution of human-specific traits (McLean et al., 2011).

### VIPs as intrinsic protein constituents of liquid-liquid phase-separated condensates contributing to fitness and adaptation of human cells

Present analyses argue that genes encoding VIPs may represent one of the principal genomic elements of human-specific regulatory networks. However, multiple lines of experimental evidence and theoretical considerations suggest that VIPs may constitute an important regulatory component of the human proteome as well by making a vital contribution to the formation and function of multimolecular complexed based on multivalent protein interactions. Intracellular biochemical reactions are spatially compartmentalized within membraneless organelles called cellular bodies, which were originally defined as nuclear or cytoplasmic non-membrane-bound structures that can be observed as independent domains using transmission electron microscopy without the requirement for antibody labelling for their visualization (Spector, 2006). Typically, these membraneless organelles represent liquid-liquid phase-separated condensates, formation of which is mediated by cooperative interactions of multivalent proteins and nucleic acid molecules (Brangwynne et al., 2009; Banjade et al., 2015; Bergeron-Sandoval et al., 2016; Banani et al., 2017). Cooperative interactions between multivalent proteins and nucleic acid molecules located in close spatial proximity within a cell initiate the formation of liquid-liquid phase-separated condensates appearing as membraneless intracellular organelles, which are being assembled to compartmentalize and accelerate specific biochemical reactions. Most recently, a phase separation model of transcriptional control in mammalian cells has been formulated (Hnisz et al., 2017) and direct experimental evidence supporting the model have been reported (Sabari et al., 2018; Cho et al., 2018).

One of the essential structural features of protein constituents of phase-separated condensates is the propensity for multivalent intermolecular interactions. Notably, it has been reported that human virus-interacting proteins (VIPs) have more interacting protein partners in the network of human protein–protein interactions than non-VIPs (Luisi et al., 2015; Stark et al., 2011; Enard and Petrov, 2018). These observations indicate that human VIPs may represent intrinsic constituents of many phase-separated membraneless intracellular organelles and these VIP-containing liquid-liquid phase-separated condensates might be pathogenic targets of viruses. Several independent lines of experimental evidence and theoretical considerations stemming from the concept of intrinsically disordered proteins (IDPs) support this hypothesis.

It has been underscored that IDPs are abundant in the mammalian and viral proteomes (Tarakhovsky and Prinjha, 2018). One of the characteristic features of IDPs is the flexibility of three-dimensional conformations and this lack of structural constraints of both host and viral IDPs is likely to promote a multiplicity of viral protein interactions with host proteins, thus targeting multiple elements of the host cell defense system (Xue et al., 2014). IDPs are unable to fold spontaneously into stable three-dimensional globular structures and continually fluctuate through a range of conformations (Dyson and Wright, 2005) because they contain high proportions of charged hydrophilic amino acids combined with a relatively low abundance of hydrophobic amino acids. This flexibility of conformational features allows IDPs to form dynamically heterogeneous complexes with multiple binding partners (Ferreon et al., 2009; Ishiyama et al., 2010). The dynamic conformational nature of IDPs is essential for increased exposure and availability of potential binding sites to multiple binding partners, thus facilitating the multivalent intermolecular interactions of IDPs within diverse sets of multiprotein complexes (Tarakhovsky and Prinjha, 2018). Collectively, these features contribute to the apparent governing role of IDPs in signaling networks (Dunker et al., 2005; Dyson and Wright, 2005; Ishiyama et al., 2010). It has been suggested that viral proteins interfere with host phase-separated protein condensates to maximize the impact of viruses on host protein networks and that both viral and host IDPs play a central role in these processes (Tarakhovsky and Prinjha, 2018). It is tempting to speculate that continuing interactions between myriads of viral and host IDPs might represent the major force of the evolutionary adaptation on molecular, cellular, and organismal levels.

Taken together, the above considerations strongly argue that human VIPs are components of a large family of IDPs that function as intrinsic protein constituents of many liquid-liquid phase-separated multimolecular complexes facilitating compartmentalization and acceleration of biochemical reactions in human cells. Consistent with this hypothesis, *BRD4*, *MED1*, and *CDK9* genes that were recently identified as the essential structural-functional elements of phase-separated condensates exerting transcriptional controls at super-enhancers (Sabari et al., 2018; Cho et al., 2018) also have been described as VIPs (see database of VIP-encoding genes reported in Enard and Petrov, 2018). Therefore, VIP-containing phase-separated multimolecular complexes are likely to represent high-value targets for viruses and important fitness determinants during adaptation.

It has been estimated that interactions with viruses accounted for ∼30% of protein adaptation in the human lineage (Enard et al., 2016; 2018), consistent with the hypothesis that viruses appear to be a major driver of adaptation in the human lineage. Virus-interacting proteins (VIPs) evolve under both stronger purifying selection and tend to adapt at much higher rates compared to similar proteins that do not interact with viruses (Enard et al., 2016; 2018). Most recent assessments of the enrichment within VIPs of introgressed segments of genomic DNA after interbreeding of Neanderthals and Modern Humans provide strong evidence that viruses drive adaptive introgression between species and that adaptive introgression had a substantial impact at VIPs after interbreeding (Enard and Petrov, 2018). This contribution extends the concept of critical importance of VIPs for evolution of Modern Humans by demonstrating that 95.9% of all human genes encoding VIPs are components of human-specific genomic regulatory networks that appear to operate in distinct types of human cells ranging from preimplantation embryos to adult dorsolateral prefrontal cortex.

### Conclusion

Observations reported in this contribution support the conclusion that Modern Humans captured unique combinations of human-specific regulatory loci, divergent subsets of which were created within genomic regions highly conserved in distinct species of six NHP separated by 30 million years of evolution. Concurrently, this unique-to-human mosaic of genomic regulatory pathways built on DNA sequences inherited from ECAs was supplemented with 12,486 created de novo HSRS. Collectively, present findings suggest that incremental genomic divergence of the human lineage has been continued throughout the primate’s evolution concurrently with the emergence and segregation of other non-human Great Ape species. This complex continuous process of genomic divergence was gradually driving speciation of *H. sapiens*, in part, by capturing and retaining the unique mosaic of genomic signatures of ECAs. Genes encoding human VIPs are emerging as principal genomic elements of this continuing evolutionary process, contributing to the molecular, cellular, and organismal adaptations via interactions with myriads of viruses.

## Methods

### Data source

#### Candidate human-specific regulatory sequences and African Apes-specific retroviral insertions

A total of 51,835 candidate HSRS and all currently known 504 insertion sites of the African Apes-specific PtERV1 retrovirus (Kronenberg et al., 2018) were analyzed in this study, detailed descriptions of which and corresponding references of primary original contributions are reported in the Tables 1-4 and Supplemental Tables S1-S12.

#### Additional Data Sources and Analytical Protocols

Solely publicly available datasets and resources were used in this contribution as well as methodological approaches and a computational pipeline validated for discovery of primate-specific gene and human-specific regulatory loci (Tay et al., 2009; Kent, 2002; Schwartz et al., 2003; Capra et al., 2013; Marnetto et al., 2014; Glinsky, 2015-2018; Guffanti et al., 2018). The analysis is based on the University of California Santa Cruz (UCSC) LiftOver conversion of the coordinates of human blocks to corresponding non-human genomes using chain files of pre-computed whole-genome BLASTZ alignments with a minMatch of 0.95 and other search parameters in default setting (http://genome.ucsc.edu/cgi-bin/hgLiftOver). Extraction of BLASTZ alignments by the LiftOver algorithm for a human query generates a LiftOver output “Deleted in new”, which indicates that a human sequence does not intersect with any chains in a given non-human genome. This indicates the absence of the query sequence in the subject genome and was used to infer the presence or absence of the human sequence in the non-human reference genome. Human-specific regulatory sequences were manually curated to validate their identities and genomic features using a BLAST algorithm and the latest releases of the corresponding reference genome databases for time periods between April, 2013 and September, 2018.

The significance of the differences in the expected and observed numbers of events was calculated using two-tailed Fisher’s exact test. Additional placement enrichment tests were performed for individual classes of HSRS taking into account the size in bp of corresponding genomic regions. Datasets of NANOG-, POU5F1-, and CTCF-binding sites and human-specific TFBS in hESCs as well as all other classes of HSRS were reported previously (Kunarso et al., 2010; McLean et al., 2011; Prüfer et al., 2012; Shulha et al., 2012; Konopka et al., 2012; Scally et al., 2012; Capra et al., 2013; Marchetto et al., 2013; Marnetto et al., 2014; Prescott et al., 2015; Gittelman et al. 2015; Glinsky et al., 2015-2018; Dong et al., 2016; Sousa et al., 2017; Dennis et al., 2017; Kronenberg et al., 2018; Guffanti et al., 2018) and are publicly available.

### Data analysis

#### Categories of DNA sequence conservation

Identification of highly-conserved in primates (pan-primate), primate-specific, and human-specific sequences was performed as previously described (Glinsky, 2015-2018). In brief, all categories were defined by direct and reciprocal mapping using liftOver (see above). Specifically:

- <u>Highly conserved in primates’ sequences</u>: DNA sequences that have at least 95% of bases remapped during conversion from/to human (Homo sapiens, hg38), chimp (Pan troglodytes, v5), and bonobo (Pan paniscus, v2; in specifically designated instances, Pan paniscus, v1 was utilized for comparisons). Similarly, highly-conserved sequences were defined for hg38 and genomes of Gorilla, Orangutan, Gibbon, and Rhesus.
- <u>Primate-specific</u>: DNA sequences that failed to map to the mouse genome (mm10).
- <u>Human-specific</u>: DNA sequences that failed to map at least 10% of bases from human to both chimpanzee and bonobo. All candidate HSRS identified based on the sequence alignments failures to genomes of both chimpanzee and bonobo were subjected to more stringent additional analyses requiring the mapping failures to genomes of Gorilla, Orangutan, Gibbon, and Rhesus. These loci were considered created de novo human-specific regulatory sequences (HSRS).

To infer the putative evolutionary origins, each evolutionary classification was defined independently by running the corresponding analyses on all candidate HSRS representing the specific category. For example, human-rodent conversion identify sequences that are absent in the mouse genome based on the sequence identity threshold of 10%). Additional comparisons were performed using the same methodology and exactly as stated in the manuscript text and described in details below.

#### Genome-wide proximity placement analysis

Genome-wide Proximity Placement Analysis (GPPA) of distinct genomic features co-localizing with HSRS was carried out as described previously (Glinsky, 2015-2018). Briefly, as typical example of the analytical protocol, we examined the significance of overlaps between hESC active enhances and hsTFBS by first identifying all hsTFBS that overlap with any of the genomic regions tested in the ChIP-STARR-seq dataset (Barakat etl, 2018; Glinsky et al., 2018). We then calculated the relative frequency of active enhancers overlapping with hsTFBS. To assess the significance of the observed overlap of genomic coordinates, we compared the values recorded for hsTFBS with the expected frequency of active and non-active enhancers that overlap with all TFBS for NANOG (15%) and OCT4 (25%) as previously determined (Barakat et al 2018). Our analyses demonstrate that more than 95% of hsTFBS co-localized with sequences in the tested regions of the hESC genome.

#### Inference of the volutionary origin and functional enrichment analyses

Evolutionary origins of HSRS were inferred from the results of the conservation patterns of 59,732 candidate human-specific regulatory DNA sequences based on the hg38 release of the human reference genome and latest available releases of genomes of six non-human primates, namely Chimpanzee, Bonobo, Gorilla, Orangutan, Gibbon, and Rhesus. The conservation analyses was carried-out using the LiftOver algorithm and Multiz Alignments of 20 mammals (17 primates) of the UCSC Genome Browser (Kent et al., 2002) on Human Dec. 2013 Assembly (GRCh38/hg38) (http://genome.ucsc.edu/cgi-bin/hgTracks?db=hg38&position=chr1%3A90820922-90821071&hgsid=441235989_eelAivpkubSY2AxzLhSXKL5ut7TN).

All DNA sequences were converted to most recent releases of the corresponding reference genome databases and were utilized consistently throughout the study to ensure the use of the most precise, accurate, and reproducible genomic DNA sequences available to date. A candidate HSRS was considered conserved if it could be aligned from/to hg38 reference genome and either one or both *Chimpanzee* or *Bonobo* genomes using defined sequence conservation thresholds of the LiftOver algorithm MinMatch function and direct and reciprocal conversions protocols. Similarly, the conservation patterns were evaluated for genomes of other NHP. LiftOver conversion of the coordinates of human blocks to non-human genomes using chain files of pre-computed whole-genome BLASTZ alignments with a specified MinMatch levels and other search parameters in default setting (http://genome.ucsc.edu/cgi-bin/hgLiftOver). Several thresholds of the LiftOver algorithm MinMatch function (minimum ratio of bases that must remap) were utilized to assess the sequences conservation and identify candidate human-specific (MinMatch of 0.1; 0.95; 0.99; and 1.00) and conserved in nonhuman primates (MinMatch of 0.95 and 1.00) regulatory sequences as previously described (Glinsky, 2015-2018; Guffanti et al., 2018). The Net alignments provided by the UCSC Genome Browser were utilized to compare the sequences in the human genome (hg38) with the mouse (mm10), *Chimpanzee* (PanTro5), and latest available releases of *Bonobo*, Gorilla, *Orangutan*, *Gibbon*, and *Rhesus* genomes. A given regulatory DNA segment was defined as the highly conserved regulatory sequence when both direct and reciprocal conversions between humans’ and nonhuman primates’ genomes were observed using the MinMatch sequence alignment threshold of 0.95 requiring that 95% of bases must remap during the alignments of the corresponding sequences. A given regulatory DNA segment was defined as the created de novo candidate human-specific regulatory sequence when sequence alignments failed to both *Chimpanzee* and *Bonobo* genomes using the specified MinMatch sequence alignment thresholds. More stringently, these requirements were extended to include genomes of Gorilla, Orangutan, Gibbon, and Rhesus. Analyses of conservation patterns of 11,866 human-specific insertions have been performed using eleven different window sizes (Supplemental Table S12) centered at the insertion sites previously reported by Kronenberg et al. (2018). Numbers of records that successfully completed direct and reciprocal conversions from/to hg38 and genomes of non-human species (six non-human primates and mouse) using sequence identity threshold 95% are reported in the Supplemental Table S12.

The Enrichr API (January 2018 version) (Chen et al., 2013) was used to test genes linked to HSRS of interest for significant enrichment in numerous functional categories. To comply with the web interface, we considered the 1000 genes closest to the tested peaks for enrichments. In all plots, we report the “combined score” calculated by Enrichr, which is a product of the significance estimate and the magnitude of enrichment (combined score *c = log(p) * z*, where *p* is the Fisher’s exact test p-value and *z* is the z-score deviation from the expected rank). Additional functional enrichment analyses were performed with GREAT (McLean et al., 2010).

##### Statistical Analyses of the Publicly Available Datasets

All statistical analyses of the publicly available genomic datasets, including error rate estimates, background and technical noise measurements and filtering, feature peak calling, feature selection, assignments of genomic coordinates to the corresponding builds of the reference human genome, and data visualization, were performed exactly as reported in the original publications and associated references linked to the corresponding data visualization tracks (http://genome.ucsc.edu/). Any modifications or new elements of statistical analyses are described in the corresponding sections of the Results. Statistical significance of the Pearson correlation coefficients was determined using GraphPad Prism version 6.00 software. Both nominal and Bonferroni adjusted p values were estimated. The significance of the differences in the numbers of events between the groups was calculated using two-sided Fisher’s exact and Chi-square test, and the significance of the overlap between the events was determined using the hypergeometric distribution test (Tavazoie et al., 1999).

## Supplemental Information

Supplemental information includes Supplemental Tables S1-S12, Supplemental Text, and Supplemental Figure S1.

## Author Contributions

This is a single author contribution. All elements of this work, including the conception of ideas, formulation, and development of concepts, execution of experiments, analysis of data, and writing of the paper, were performed by the author.

## Acknowledgements

This work was made possible by the open public access policies of major grant funding agencies and international genomic databases and the willingness of many investigators worldwide to share their primary research data. I would like to thank my anonymous colleagues for their valuable critical contributions during the peer review process of this work.

