## Supplemental Text for "Analysis of genomic loci harboring 59,732 human-specific regulatory sequences reveals unique to human regulatory patterns associated with brain development"

**Insertion sites of the African Great Ape-specific retrovirus PtERV1 and significant fractions of distinct classes HSRS share common genomic coordinates**

Structurally distinct mutations within genomic regions harboring HSRS that independently emerged on the Modern Humans lineage and distinct species of non-human Great Apes are of particular interest because they might indicate the functional divergence between species of these independently-targeted regulatory regions. In this context, it was of interest to determine whether genomic regions harboring HSRS intersect genomic coordinates of insertion sites of the African Great Ape-specific retrovirus PtERV1. Significantly, no PtERV1 insertions were detected in genomes of Modern Humans and Orangutan yet the PtERV1 retrovirus appears integrated at 540 loci in genomes of Gorilla, Chimpanzee, and Bonobo during millions years of evolution (Kronenberg et al., 2018). Analysis of evolutionary patterns of insertions of African ape-specific retrovirus PtERV1 revealed that, without exception, all distinct classes of HSRS analyzed in this study intersect within 10 Kb windows of orthologous genomic regions targeted by PtERV1 in genomes of Gorilla, Chimpanzee, and Bonobo, albeit with different degrees of frequencies and significance (Supplemental Tables S6a; S6b; S6c). Interestingly, genomic coordinates of PtERV1 insertions in the Gorilla genome appear to overlap more frequently genomic regions harboring HSRS (Supplemental Table S6b).

Genome-wide analysis of gene expression changes in human-chimpanzee cerebral organoids revealed that human-specific duplications, in contrast to other types of human-specific structural variations, are associated with up-regulated genes in human radial glial and excitatory neurons (Kronenberg et al., 2018). These observations are highly consistent with previous studies demonstrating that defined human-specific segmental duplications of *SRGAP2* and *ARHGAP11B* genes drive phenotypic differences in cortical development between humans and chimpanzee (Dennis et al., 2012; Charrier et al., 2012; Florio et al., 2015). Proximity placement enrichment analysis of 7,897 duplication regions in human genome and PtERV1 insertion sites (Supplemental Table S6c) identified 71 PtERV1 loci intersecting 87 duplication regions and revealed significantly more frequent co-localization of Chimpanzee-specific PtERV1 insertions compared to Gorilla-specific insertions (24.2% versus 10.7%, respectively; p = 0.0076; 2-tailed Fisher’s exact test). Overall, these analyses identified 248 PtERV1 loci (45.9%; p = 1.03E-44; hypergeometric distribution test) intersecting human genomic regions harboring 442 candidate HSRS (Supplemental Tables S6a; S6b; and S6c), which are significantly enriched (p = 0.0018) for regions of fixed human-specific mutations that have been associated with human-specific changes of gene expression in cerebral organoids’ models of brain development (Kronenberg et al., 2018). This set of genomic regions with overlapping coordinates of PtERV1 integration sites and loci harboring human-specific mutations of potential functional significance may represent an attractive functional validation panel of elite candidate regulatory sequences likely contributing to phenotypic divergence of Modern Humans and our closest evolutionary relatives.
