## Supplemental Figure 1 for "Analysis of genomic loci harboring 59,732 human-specific regulatory sequences reveals unique to human regulatory patterns associated with brain development"

### Slide 1
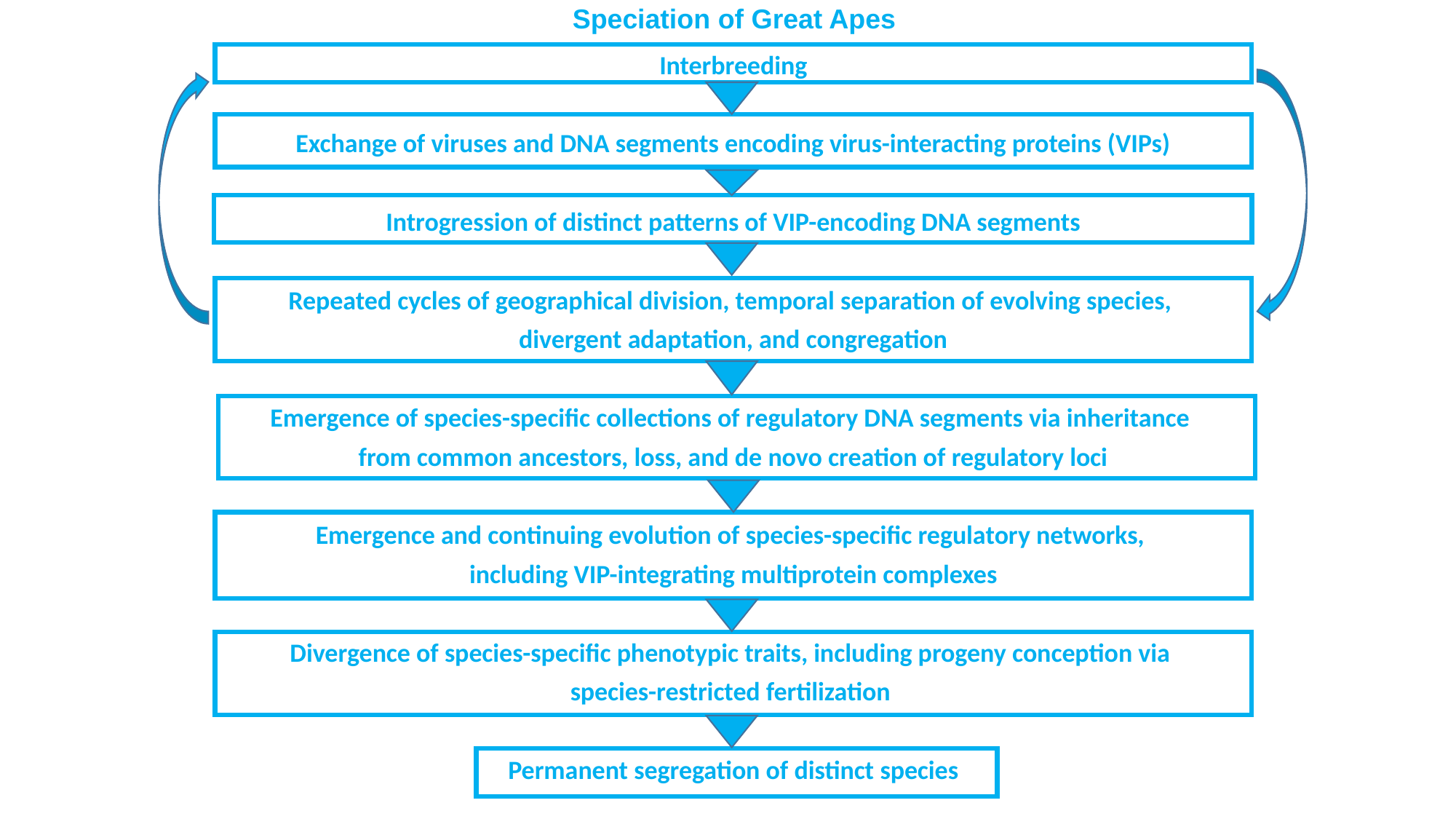

#### Speciation of Great Apes
Interbreeding
Exchange of viruses and DNA segments encoding virus-interacting proteins (VIPs)
Introgression of distinct patterns of VIP-encoding DNA segments
Repeated cycles of geographical division, temporal separation of evolving species,
divergent adaptation, and congregation
Emergence of species-specific collections of regulatory DNA segments via inheritance
from common ancestors, loss, and de novo creation of regulatory loci
Emergence and continuing evolution of species-specific regulatory networks,
including VIP-integrating multiprotein complexes
Divergence of species-specific phenotypic traits, including progeny conception via
species-restricted fertilization
Permanent segregation of distinct species

### Slide 2
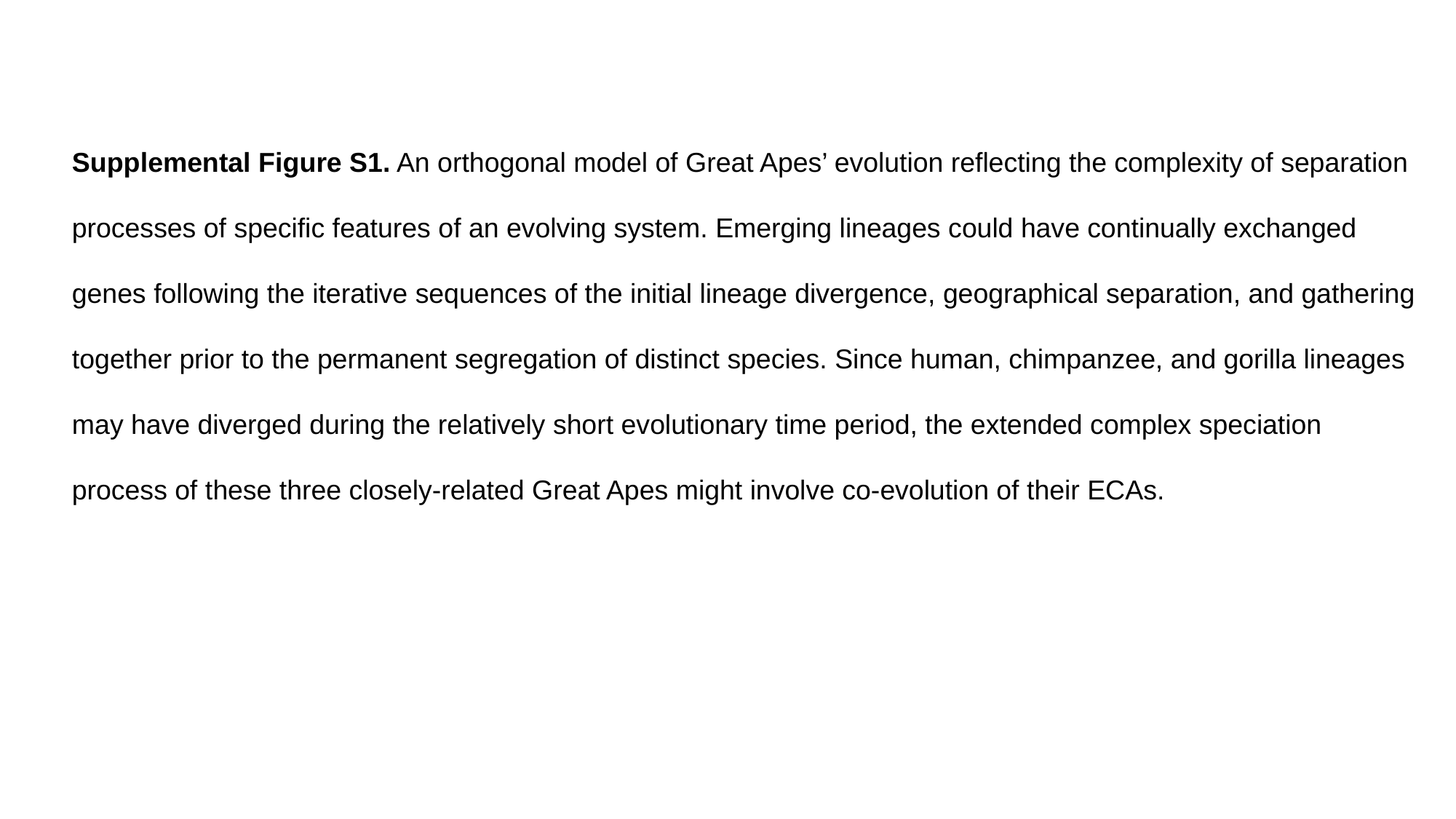

Supplemental Figure S1. An orthogonal model of Great Apes’ evolution reflecting the complexity of separation processes of specific features of an evolving system. Emerging lineages could have continually exchanged genes following the iterative sequences of the initial lineage divergence, geographical separation, and gathering together prior to the permanent segregation of distinct species. Since human, chimpanzee, and gorilla lineages may have diverged during the relatively short evolutionary time period, the extended complex speciation process of these three closely-related Great Apes might involve co-evolution of their ECAs.
