## Supplemental Tables S1-S5; S11; S12 for "Analysis of genomic loci harboring 59,732 human-specific regulatory sequences reveals unique to human regulatory patterns associated with brain development"

**Supplemental Table S1.** A catalog of human-specific genomic regulatory loci and networks reported to date.

| **Candidate human-specific regulatory loci** | **References** |
|---|---|
| Regions of human-specific loss of conserved regulatory DNA termed hCONDEL | McLean et al., 2011 |
| Human-specific epigenetic regulatory marks consisting of H3K4me3 histone methylation signatures at transcription start sites in prefrontal neurons | Shulha et al., 2012 |
| Human-specific transcriptional genetic networks in the frontal lobe | Konopka et al., 2012 |
| Conserved in humans regulatory DNA sequences designated human accelerated regions, HARs | Capra et al., 2013 |
| Fixed human-specific regulatory regions, FHSRR | Marnetto et al., 2014 |
| Human-specific transcription factor-binding sites, HSTFBS, and hESC enhancers | Glinsky, 2015-2018 |
| DNase I hypersensitive sites (DHSs) that are conserved in non-human primates but accelerated in the human lineage | Gittelman et al. 2015 |
| DNase I hypersensitive sites (DHSs) that are under accelerated evolution, ace-DHSs | Dong et al., 2016 |
| Fixed human-specific insertions | Kronenberg et al., 2018 |
| Fixed human-specific deletions | Kronenberg et al., 2018 |
| Human-specific short tandem repeats (STR) expansions | Kronenberg et al., 2018 |
| Human-specific short tandem repeats (STR) contractions | Kronenberg et al., 2018 |
| Human-specific mutations associated with human-specific gene expression changes in excitatory neurons | Kronenberg et al., 2018 |
| Human-specific mutations associated with human-specific gene expression changes in radial glia | Kronenberg et al., 2018 |

| | |
|---|---|
| Duplicated regions in GRCh38 space defined by WSSD | Kronenberg et al., 2018 |
| Human-specific hESC functional enhancers | Glinsky et al., 2018 |
| Human-specific gene expression signatures of induced pluripotent stem cells | Marchetto et al., 2013 |
| Human-specific segmental duplications | Dennis MY et al., 2017 |
| Genes with firmly established neurodevelopmental functions and well-documented genetic/genomic/epigenetic alterations of potential functional significance acquired within the human lineage after the divergence of humans and chimpanzees | Sousa, et al. 2017 |
| Chimp-biased developmental enhancers | Prescott et al., 2015 |
| Human-biased developmental enhancers | Prescott et al., 2015 |
| Human-specific TE loci expressed in DLPFC | Guffanti et al., 2018 |

**Supplemental Table S2.** Human-specific regulatory sequences (HSRS) analyzed in this contribution

| Human-specific regulatory sequences (HSRS) | Number of loci | References |
|---|---|---|
| Human-specific STR contractions | 1464 | Kronenberg et al., 2018 |
| Human-specific STR expansions | 4910 | Kronenberg et al., 2018 |
| Fixed human-specific deletions | 5891 | Kronenberg et al., 2018 |
| Fixed human-specific insertions | 11886 | Kronenberg et al., 2018 |
| All regions of human-specific mutations | 24151 | Kronenberg et al., 2018 |
| Duplicated regions in GRCh38 space defined by WSSD | 7897 | Kronenberg et al., 2018 |
| Fixed human-specific regulatory regions (FHSRR) | 4249 | Marnetto et al., 2014 |
| Accelerated evolution-DHS (ace-DHS) | 3538 | Dong et al., 2016 |
| Human accelerated regions (HARs) | 2741 | Capra et al., 2013 |
| Chimp-biased developmental enhancers | 999 | Prescott et al., 2015 |
| Human-specific segmental duplications | 218 | Dennis et al., 2017 |
| Human-biased developmental enhancers | 996 | Prescott et al., 2015 |
| DHS Fixed human-specific regulatory regions (DHS-FHSRR) | 2116 | Marnetto et al., 2014 |
| Human-specific functional enhancers in hESC | 1619 | Glinsky et al., 2018; Glinsky, 2018; Barakat et al., 2018 |
| Human accelerated DHS (haDHS) | 524 | Gittelman et al. 2015 |
| Human-specific CTCF binding sites in hESC | 575 | Glinsky, 2015-2016; Kunarso et al., 2010 |

| | | |
|---|---|---|
| Human-specific OCT4 binding sites in hESC | 2328 | Glinsky, 2015-2016; Kunarso et al., 2010 |
| Human-specific H3K4me3 peaks in prefrontal cortex | 406 | Shulha et al., 2012 |
| Human-specific NANOG binding sites in hESC | 816 | Glinsky, 2015-2016; Kunarso et al., 2010 |
| hESC fixed human-specific regulatory regions (hESC-FHSRR) | 1932 | Marnetto et al., 2014 |
| Human-specific TE loci expressed in DLPFC | 4627 | Guffanti et al, 2018; Glinsky, 2018 (this work) |
| Total candidate HSRS | 59,732 | |

**Supplemental Table S3.** Human-specific regulatory sequences (HSRS) associated with human-specific gene expression changes in the cerebral organoid model of brain development

| | |
|---|---|
| HSRS associated with human-specific gene expression in cerebral organoids | 1466 |
| HSRS associated with human-specific gene expression changes in Radial glia | 947 |
| HSRS associated with human-specific gene expression changes in Excitatory neurons | 517 |
| HSRS associated with genes Down-regulated in Radial glia of the human developing brain | 417 |
| HSRS associated with genes Up-regulated in Radial glia of the human developing brain | 531 |
| HSRS associated with genes Down-regulated in Excitatory neurons of the human developing brain | 227 |
| HSRS associated with genes Up-regulated in Excitatory neurons of the human developing brain | 291 |

Kronenberg et al., 2018

High-resolution comparative analysis of great ape genomes. Kronenberg et al., Science 360, 1085 (2018)

**Supplemental Table S4.** Mosaicism of evolutionary origins of 7,897 duplicated regions in GRCh38 space defined by whole-genome shotgun sequence detection (WSSD)

| Non-human species (NHS) | Number of highly conserved regions converted from/to hg38 | Percent | Loss of ancestral loci | Percent | Non-human species | Number of highly conserved regions converted from/to hg38 | Percent |
|---|---|---|---|---|---|---|---|
| Gorilla all records | 5571 | 81.61 | 1255 | 18.39 | Gorilla only records | 458 | 6.71 |
| Chimpanzee all records | 5484 | 80.34 | 1342 | 19.66 | Chimpanzee only records | 454 | 6.65 |
| Bonobo all records | 4862 | 71.23 | 1964 | 28.77 | Bonobo only records | 362 | 5.30 |
| Orangutan all records | 3744 | 54.85 | 3082 | 45.15 | Orangutan only records | 313 | 4.59 |
| Gibbon all records | 3331 | 48.80 | 3495 | 51.20 | Gibbon only records | 281 | 4.12 |
| Rhesus all records | 2951 | 43.23 | 3875 | 56.77 | Rhesus only records | 251 | 3.68 |
| Mouse all records | 451 | 6.61 | 6375 | 93.39 | Mouse only records | 58 | 0.85 |
| NHS all highly-conserved records | 6826 | 100.00 | 0 | 0.00 | NHS all species-specific highly-conserved records | 2177 | 31.89 |

Legend: reported only individual highly-conserved regions uniquely remapped to hg38 during reciprocal conversions from corresponding NHS genomes;

**Supplemental Table S5.** Mosaicism of evolutionary origins of genomic loci harboring candidate human-specific regulatory sequences associated with human-specific gene expression changes in human versus chimpanzee brain organoids

| **Classification category** | Human genome | Mapped to highly | Percent | Chimpanzee | Percent | Bonobo | Percent | Gorilla | Percent | Orangutan | Percent | Gibbon | Percent | Rhesus | Percent | Mouse | Percent |
|---|---|---|---|---|---|---|---|---|---|---|---|---|---|---|---|---|---|
| **Human-specific mutations associated with human-specific gene expression changes in excitatory neurons**** | 517 | 283 | 54.74 | **183** | 64.66 | **143** | 50.53 | **249** | 87.99 | **9** | 3.18 | 16 | 5.65 | **14** | 4.95 | 0 | 0.00 |
| **Human-specific mutations associated with human-specific gene expression changes in radial glia**** | 947 | 526 | 55.54 | **326** | 61.98 | **282** | 53.61 | **472** | 89.73 | **21** | 3.99 | 24 | 4.56 | **27** | 5.13 | 0 | 0.00 |

Only records remapped to the identical loci; **Associated with changes in gene expression in human versus chimpanzee brain organoids;

**Supplemental Table S11.** Enrichment within human-specific regulatory networks of human virus-interacting proteins (VIPs)

**Networks of genes associated with expression of transposable elements (TE) in human dorsolateral prefrontal cortex (DLPFC)**

| Classification category | Number of genes | Human VIPs |
|---|---|---|
| Human genome | 63677 | 4433 |
| Networks of genes associated with human DLPFC-expressed TE | 22863 | 3574 |
| Percent | 35.90 | 80.62 |
| Enrichment** | 1.00 | 2.25 |
| P value* | | 0 |

**GES of the Multilineage Markers Expressing (MLME) cells of human preimplantation embryo**

| Classification category | Number of genes | Human VIPs |
|---|---|---|
| Human genome | 63677 | 4433 |
| GES of the MLME cells of human preimplantation embryo | 12735 | 3551 |
| Percent | 20.00 | 80.10 |
| Enrichment** | 1.00 | 4.01 |
| P value* | | 0 |

**Regulatory networks of genes associated with human-specific structural variants*****

| Classification category | Number of genes | Number of human VIPs |
|---|---|---|
| Human genome | 63677 | 4433 |
| Genes associated with human-specific deletions and insertions | 10992 | 1207 |
| Percent | 17.26 | 27.23 |
| Enrichment** | 1.00 | 1.58 |
| P value* | | 1.03E-66 |

**HERVH/LBP9 pathway in hESC**

| Classification category | Number of genes | Number of human VIPs |
|---|---|---|
| Human genome | 63677 | 4433 |
| Genes associated with the HERVH/LBP9 pathway in hESC | 11507 | 2661 |
| Percent | 18.07 | 60.03 |
| Enrichment** | 1.00 | 3.32 |
| P value* | | 0 |

Legend: *, p values were estimate using the hypergeometric distribution test; **, expected values were estimated based on the number of genes in the human genome (63,677) and the number of genes in the corresponding category of human-specific regulatory networks; ***, this category of genes was reported in Kronenberg et al. (2018); TE, transposable genetic elements; hESC, human embryonic stem cells; DLPFC, dorsolateral prefrontal cortex; MLME, multi lineage markers expression; Total number of human 4,433 genes (hg38) encoding VIPs was obtained after the adjustment for loci having multiple ENSEMBL IDs associated with same Gene Symbols and removal of 3 genes that are no longer in the ENSEMBL database.

**Supplemental Table S12.** Conservation patterns of genomic regions harboring 11,866 fixed human-specific insertions**.**

| Region size/Species | Chimpanzee* | Percent | Bonobo | Percent | Gorilla | Percent | Orangutan | Percent | Rhesus | Percent | Mouse | Percent |
|---|---|---|---|---|---|---|---|---|---|---|---|---|
| 50 bp | 100 | 0.84 | 130 | 1.10 | 111 | 0.94 | 103 | 0.87 | 128 | 1.08 | 92 | 0.78 |
| 100 bp | 50 | 0.42 | 73 | 0.62 | 50 | 0.42 | 63 | 0.53 | 66 | 0.56 | 25 | 0.21 |
| 200 bp | 11 | 0.09 | 14 | 0.12 | 40 | 0.34 | 30 | 0.25 | 55 | 0.46 | 4 | 0.03 |
| 500 bp | 4 | 0.03 | 24 | 0.20 | 3 | 0.03 | 17 | 0.14 | 41 | 0.35 | 1 | 0.01 |
| 1100 bp | 53 | 0.45 | 63 | 0.53 | 49 | 0.41 | 28 | 0.24 | 52 | 0.44 | 0 | 0.00 |
| 2000 bp | 415 | 3.50 | 409 | 3.45 | 398 | 3.35 | 276 | 2.33 | 215 | 1.81 | 0 | 0.00 |
| 4000 bp | 1165 | 9.82 | 1057 | 8.91 | 1119 | 9.43 | 626 | 5.28 | 429 | 3.62 | 0 | 0.00 |
| 6000 bp | 1897 | 15.99 | 1665 | 14.03 | 1822 | 15.35 | 886 | 7.47 | 536 | 4.52 | 0 | 0.00 |
| 8000 bp | 6912 | 58.25 | 6036 | 50.87 | 6765 | 57.01 | 3061 | 25.80 | 1312 | 11.06 | 0 | 0.00 |
| 10000 bp | 7138 | 60.16 | 6261 | 52.76 | 7104 | 59.87 | 3209 | 27.04 | 2036 | 17.16 | 0 | 0.00 |

Legend: *Number of records that successfully completed direct and reciprocal conversions from/to hg38 and genomes of non-human primates and mouse using sequence identity threshold 95% of genomic regions centered at the insertion sites
