## Supplemental Table S6a for "Analysis of genomic loci harboring 59,732 human-specific regulatory sequences reveals unique to human regulatory patterns associated with brain development"

**Supplemental Table S6a.** Distribution profiles of human genomic regions harboring 442 loci of distinct classes of candidate HSRS co-localizing with insertion sites of 248 PtERV1 loci in Chimpanzee, Gorilla, and Bonobo genomes.

| Human-specific regulatory sequences (HSRS) | Number of loci | Intersecting PtERV1 loci | Intersecting human-specific loci | Percent | P value** |
| --- | --- | --- | --- | --- | --- |
| Human-specific STR contractions* | 1464 | 4 | 4 | 0.27 |  |
| Human-specific STR expansions | 4910 | 25 | 25 | 0.51 | 0.374 |
| Fixed human-specific deletions | 5891 | 38 | 37 | 0.63 | 0.118 |
| Fixed human-specific insertions | 11886 | 66 | 66 | 0.56 | 0.182 |
| All regions of human-specific mutations | 24151 | 133 | 132 | 0.55 | 0.195 |
| Fixed human-specific regulatory regions (FHSRR) | 4249 | 15 | 31 | 0.73 | 0.053 |
| Accelerated evolution-DHS (ace-DHS) | 3538 | 23 | 25 | 0.71 | 0.068 |
| Human accelerated regions (HARs) | 2741 | 12 | 11 | 0.40 | 0.597 |
| Chimp-biased developmental enhancers | 999 | 5 | 5 | 0.50 | 0.499 |
| Human segmental duplications | 218 | 2 | 2 | 0.92 | 0.176 |
| Duplicated regions in GRCh38 space defined by WSSD | 7897 | 71 | 87 | 1.10 | 0.001 |
| Human-biased developmental enhancers | 996 | 9 | 10 | 1.00 | 0.026 |
| DHS Fixed human-specific regulatory regions (DHS-FHSRR)* | 2116 | 4 | 17 | 0.80 | 0.046 |
| Human-specific functional enhancers in hESC | 1619 | 10 | 14 | 0.86 | 0.034 |
| Human accelerated DHS (haDHS)* | 524 | 1 | 1 | 0.19 | 1 |
| Human-specific CTCF binding sites in hESC* | 575 | 2 | 3 | 0.52 | 0.410 |
| Human-specific OCT4 binding sites in hESC* | 2328 | 3 | 4 | 0.17 | 0.495 |
| Human-specific H3K4me3 peaks in prefrontal cortex | 406 | 5 | 5 | 1.23 | 0.027 |
| Human-specific NANOG binding sites in hESC | 816 | 14 | 19 | 2.33 | 5.04E-06 |
| hESC fixed human-specific regulatory regions (hESC-FHSRR) | 1932 | 14 | 29 | 1.50 | 0.00026 |
| Human-specific TE loci expressed in DLPFC | 4627 | 23 | 47 | 1.02 | 0.0046 |
| Human-specific gene expression in brain organoids | 1466 | 19 | 21 | 1.43 | 0.00087 |
| Radial glia | 947 | 17 | 19 | 2.01 | 3.43E-05 |
| Excitatory neurons | 517 | 5 | 5 | 0.97 | 0.0576 |
| Radial glia Down | 417 | 13 | 14 | 3.36 | 7.26E-07 |
| Radial glia Up | 531 | 4 | 5 | 0.94 | 0.062 |
| Excitatory neurons Down | 227 | 3 | 3 | 1.32 | 0.055 |
| Excitatory neurons Up | 291 | 2 | 2 | 0.69 | 0.261 |
| Total number of genomic regions harboring HSRS | 59732 | 248 | 442 | 0.74 |  |

Legend: * the overlap of genomic coordinates of PtERV1 insertions and HSRS-harboring regions of this regulatory category is not statistically significant based on the hypergeometric distribution test; ** p values were estimated using the two-tailed Fisher's exact test compared to Human-specific STR contractions category; HSRS, human-specific regulatory sequences; WSSD, whole-genome shotgun sequence detection; underlined text denotes statistically significant categories estimated by both hypergeometric distribution test and two-tailed Fisher’s exact test;
