## Supplemental Table S6b for "Analysis of genomic loci harboring 59,732 human-specific regulatory sequences reveals unique to human regulatory patterns associated with brain development"

**Supplemental Table S6b.** Evolutionary patterns of Chimpanzee, Gorilla, and Bonobo PtERV1 insertions intersecting genomic regions harboring HSRS.

| Taxa* | Number of PtERV1 loci | PtREV1 loci intersecting genomic regions harboring human-specific regulatory loci | Percent | P value** |
| --- | --- | --- | --- | --- |
| gorilla only | 280 | 111 | 39.64 | 6.41E-20 |
| chimp,gorilla | 5 | 3 | 60.00 | 0.033 |
| chimp,chimp,bonobo,gorilla,gorilla | 1 | 1 | 100.00 | 0.168 |
| chimp,chimp,bonobo,gorilla | 5 | 0 | 0.00 | 0.399 |
| chimp,chimp,bonobo | 150 | 55 | 36.67 | 2.66E-09 |
| chimp,bonobo | 17 | 8 | 47.06 | 0.0029 |
| chimp only | 66 | 21 | 31.82 | 0.0012 |
| bonobo,chimp,gorilla | 1 | 0 | 0.00 | 0.832 |
| bonobo,chimp | 2 | 1 | 50.00 | 0.279 |
| bonobo only | 13 | 3 | 23.08 | 0.215 |
| All PtERV1 insertions | 540 | 203 | 37.59 | 3.24E-31 |
| Gorilla all PtERV1 insertions | 292 | 115 | 39.38 | 2.62E-20 |
| Chimpanzee all PtERV1 insertions | 247 | 89 | 36.03 | 1.71E-13 |
| Bonobo all PtERV1 insertions | 189 | 60 | 31.75 | 1.94E-07 |

Legend: * evolutionary patterns of 540 PtERV1 loci in genomes of non-human apes were reported by Kronenberg et al. (2018); ** p values were estimated using the hypergeometric distribution test considering the number of non-overlapping 10Kb regions in the human genome (308,829); the number of analyzed regions harboring HSRS (51,835); and corresponding numbers of the PtERV1 loci; no significant differences were observed between different categories;
