## Supplemental Table S6c for "Analysis of genomic loci harboring 59,732 human-specific regulatory sequences reveals unique to human regulatory patterns associated with brain development"

**Supplemental Table S6c.** Evolutionary patterns of Chimpanzee, Gorilla, and Bonobo PtERV1 insertions intersecting duplicated regions of human genome (hg38) defined by the whole-genome shotgun sequence detection (WSSD).

| Taxa* | Number of PtERV1 loci | PtREV1 loci intersecting genomic regions harboring human-specific regulatory loci | Percent | P value** |
| --- | --- | --- | --- | --- |
| gorilla only*** | 280 | 30 | 10.71 | 4.98E-11 |
| chimp,gorilla | 5 | 1 | 20.00 | 0.115 |
| chimp,chimp,bonobo,gorilla,gorilla | 1 | 1 | 100.00 | 0.026 |
| chimp,chimp,bonobo,gorilla | 5 | 0 | 0.00 | 0.879 |
| chimp,chimp,bonobo | 150 | 18 | 12.00 | 5.64E-08 |
| chimp,bonobo | 17 | 5 | 29.41 | 4.95E-05 |
| chimp only*** | 66 | 16 | 24.24 | 7.73E-12 |
| bonobo,chimp,gorilla | 1 | 0 | 0.00 | 0.974 |
| bonobo,chimp | 2 | 0 | 0.00 | 0.950 |
| bonobo only | 13 | 0 | 0.00 | 0.714 |
| All PtERV1 insertions | 540 | 71 | 13.15 | 3.64E-29 |
| Gorilla all PtERV1 insertions | 292 | 32 | 10.96 | 6.50E-12 |
| Chimpanzee all PtERV1 insertions | 247 | 41 | 16.60 | 2.60E-21 |
| Bonobo all PtERV1 insertions | 189 | 24 | 12.70 | 1.26E-10 |

Legend: * evolutionary patterns of 540 PtERV1 loci in genomes of non-human apes were reported by Kronenberg et al. (2018); ** p values were estimated using the hypergeometric distribution test considering the number of non-overlapping 10Kb regions in the human genome (308,829); the number of analyzed duplication regions defined in human genome by WSSD (7,897); and corresponding numbers of the PtERV1 loci; no significant differences were observed between different categories; only PtERV1 loci intersecting duplication regions in hg38 are reported; *** indicated significantly different values for Gorilla only versus Chimpanzee only records (p = 0.0076; two-tailed Fisher's exact test);
