## Supplemental Table S7 for "Analysis of genomic loci harboring 59,732 human-specific regulatory sequences reveals unique to human regulatory patterns associated with brain development"

**Supplemental Table S7.** Mosaicism of evolutionary origins of candidate human-specific regulatory loci defined based on the mapping failure to both Chimpanzee and Bonobo reference genomes.*

| Classification category | Chimpanzee | Bonobo | Rhesus | Gibbon | Orangutan | Gorilla | Human |
| --- | --- | --- | --- | --- | --- | --- | --- |
| Human-specific DLPFC-expressed transposons (n)** | 0 | 0 | 1045 | 1148 | 1371 | 1688 | 4645 |
| Percent | 0.00 | 0.00 | 22.50 | 24.71 | 29.52 | 36.34 | 100.00 |
| Human-specific hESC functional enhancers (n) | 0 | 0 | 84 | 88 | 122 | 174 | 1619 |
| Percent | 0.00 | 0.00 | 5.19 | 5.44 | 7.54 | 10.75 | 100.00 |
| Fixed human-specific regulatory regions (FHSRR) (n) | 0 | 0 | 8 | 18 | 0 | 173 | 3273 |
| Percent | 0.00 | 0.00 | 0.24 | 0.55 | 0.00 | 5.29 | 100.00 |
| hESC fixed human-specific regulatory regions (hESC-FHSRR) | 0 | 0 | 2 | 2 | 0 | 70 | 1233 |
| Percent | 0.00 | 0.00 | 0.16 | 0.16 | 0.00 | 5.68 | 100.00 |
| DHS fixed human-specific regulatory regions (DHS-FHSRR) | 0 | 0 | 1 | 4 | 0 | 79 | 1104 |
| Percent | 0.00 | 0.00 | 0.09 | 0.36 | 0.00 | 7.16 | 100.00 |
| Human-specific STR expansions (n) | 0 | 0 | 35 | 47 | 30 | 16 | 1783 |
| Percent | 0.00 | 0.00 | 1.96 | 2.64 | 1.68 | 0.90 | 100.00 |
| Fixed human-specific deletions (n) | 0 | 0 | 4 | 6 | 6 | 9 | 58 |
| Percent | 0.00 | 0.00 | 6.90 | 10.34 | 10.34 | 15.52 | 100.00 |
| Fixed human-specific insertions (n) | 0 | 0 | 0 | 0 | 0 | 1 | 7 |
| Percent | 0.00 | 0.00 | 0.00 | 0.00 | 0.00 | 14.29 | 100.00 |
| Human-specific NANOG binding sites (n) | 0 | 0 | 1 | 4 | 0 | 52 | 540 |
| Percent | 0.00 | 0.00 | 0.19 | 0.74 | 0.00 | 9.63 | 100.00 |
| Human-specific OCT4 binding sites (n) | 0 | 0 | 0 | 5 | 0 | 19 | 1791 |
| Percent | 0.00 | 0.00 | 0.00 | 0.28 | 0.00 | 1.06 | 100.00 |
| Human-specific CTCF binding sites (n) | 0 | 0 | 0 | 0 | 0 | 14 | 478 |
| Percent | 0.00 | 0.00 | 0.00 | 0.00 | 0.00 | 2.93 | 100.00 |
| Human-specific STR contractions (n) | 0 | 0 | 7 | 9 | 11 | 15 | 199 |
| Percent | 0.00 | 0.00 | 3.52 | 4.52 | 5.53 | 7.54 | 100.00 |
| All human-specific loci (n) | 0 | 0 | 1187 | 1331 | 1540 | 2310 | 16730 |
| Percent | 0.00 | 0.00 | 7.10 | 7.96 | 9.21 | 13.81 | 100.00 |

Legend: *Mosaicism of evolutionary origins was defined based on the sequence conservation analyses of human-specific regulatory loci that failed conversions to both Chimpanzee and Bonobo genomes and manifested at least 95% sequence conservations during both direct and reciprocal conversions to genomes of other non-human primates (Gorilla; Orangutan; Gibbon; Rhesus); DLPFC, dorsolateral prefrontal cortex;
