## Supplemental Table S8 for "Analysis of genomic loci harboring 59,732 human-specific regulatory sequences reveals unique to human regulatory patterns associated with brain development"

**Supplemental Table S8.** Human-specific regulatory loci defined based on the conversion failure to reference genomes of Chimpanzee, Bonobo, Gorilla, Orangutan, Gibbon, and Rhesus.*

| Classification category | Human genome (hg38) | Mapping failures to both Chimpanzee and Bonobo genomes | Mapping failures to genomes of six non-human primates |
| --- | --- | --- | --- |
| Human-specific hESC functional enhancers (n) | 1619 | 1619 | 1343 |
| Percent | 100.00 | 100.00 | 82.95 |
| Human-specific CTCF binding sites in hESC (n) | 575 | 478 | 458 |
| Percent | 100.00 | 83.13 | 79.65 |
| Human-specific OCT4 binding sites in hESC (n) | 2328 | 1791 | 1731 |
| Percent | 100.00 | 76.93 | 74.36 |
| Fixed human-specific regulatory regions (FHSRR) (n) | 4249 | 3273 | 3035 |
| Percent | 100.00 | 77.03 | 71.43 |
| hESC fixed human-specific regulatory regions (hESC-FHSRR) | 1932 | 1233 | 1155 |
| Percent | 100.00 | 63.82 | 59.78 |
| Human-specific NANOG binding sites (n) | 816 | 540 | 483 |
| Percent | 100.00 | 66.18 | 59.19 |
| Human-specific transposons expressed in dorsolateral prefrontal cortex (n) | 4645 | 4612 | 2559 |
| Percent | 100.00 | 99.29 | 55.09 |
| DHS fixed human-specific regulatory regions (DHS-FHSRR) | 2116 | 1104 | 753 |
| Percent | 100.00 | 52.17 | 35.59 |
| Human-specific STR expansions | 4910 | 1783 | 692 |
| Percent | 100.00 | 36.31 | 14.09 |
| Human-specific STR contractions | 1464 | 199 | 183 |
| Percent | 100.00 | 13.59 | 12.50 |
| Fixed human-specific deletions | 5891 | 58 | 38 |
| Percent | 100.00 | 0.98 | 0.65 |
| Fixed human-specific insertions | 11886 | 7 | 4 |
| Percent | 100.00 | 0.06 | 0.03 |
| Duplicated regions in GRCh38 space defined by WSSD | 7897 | 75 | 52 |
| Percent | 100.00 | 0.95 | 0.66 |
| All candidate human-specific regulatory loci (n) | 50328 | 16697 | 12486 |
| Percent | 100.00 | 33.18 | 24.81 |

Legend: ** WSSD, whole-genome shotgun sequence detection; hESC, human embryonic stem cells; STR, short tandem repeats;
