## Supplemental Table S9 for "Analysis of genomic loci harboring 59,732 human-specific regulatory sequences reveals unique to human regulatory patterns associated with brain development"

**Supplemental Table S9.** Enrichment within human-specific regulatory networks of genes comprising expression signatures (GES) of human-specific neurodevelopmental transcriptional networks of excitatory neurons and radial glia.

**Networks of genes associated with expression of transposable elements (TE) in human dorsolateral prefrontal cortex**

| Classification category | Number of genes | Excitatory neurons | DOWN Humans vs Chimp | UP Humans vs Chimp | Radial glia | DOWN Humans vs Chimp | UP Humans vs Chimp |
| --- | --- | --- | --- | --- | --- | --- | --- |
| Human genome | 63677 | 384 | 165 | 219 | 668 | 285 | 383 |
| Networks of genes associated with human DLPFC-expressed TE | 22863 | 322 | 131 | 191 | 561 | 230 | 331 |
| Percent | 35.90 | 83.85 | 79.39 | 87.21 | 83.98 | 80.70 | 86.42 |
| Enrichment** | 1.00 | 2.34 | 2.21 | 2.43 | 2.34 | 2.25 | 2.41 |
| P value* |  | 5.57E-84 | 2.61E-30 | 5.05E-56 | 3.08E-146 | 2.04E-54 | 9.93E-94 |

**GES of the Multi-lineage Markers Expressing (MLME) cells of human preimplantation embryo**

| Classification category | Number of genes | Excitatory neurons | DOWN Humans vs Chimp | UP Humans vs Chimp | Radial glia | DOWN Humans vs Chimp | UP Humans vs Chimp |
| --- | --- | --- | --- | --- | --- | --- | --- |
| Human genome | 63677 | 384 | 165 | 219 | 668 | 285 | 383 |
| GES of the MLME cells of human embryo | 12735 | 262 | 111 | 151 | 481 | 209 | 272 |
| Percent | 20.00 | 68.23 | 67.27 | 68.95 | 72.01 | 73.33 | 71.02 |
| Enrichment** | 1.00 | 3.41 | 3.36 | 3.45 | 3.60 | 3.67 | 3.55 |
| P value* |  | 1.59E-93 | 1.50E-39 | 2.02E-55 | 3.95E-187 | 3.63E-84 | 1.22E-103 |

**Regulatory networks of genes associated with human-specific structural variants*****

| Classification category | Number of genes | Excitatory neurons | DOWN Humans vs Chimp | UP Humans vs Chimp | Radial glia | DOWN Humans vs Chimp | UP Humans vs Chimp |
| --- | --- | --- | --- | --- | --- | --- | --- |
| Human genome | 63677 | 384 | 165 | 219 | 668 | 285 | 383 |
| Genes associated with human-specific deletions and insertions | 10992 | 123 | 53 | 70 | 125 | 59 | 66 |
| Percent | 17.26 | 32.03 | 32.12 | 31.96 | 18.71 | 20.70 | 17.23 |
| Enrichment** | 1.00 | 1.86 | 1.86 | 1.85 | 1.08 | 1.20 | 1.00 |
| P value* |  | 6.75E-13 | 1.39E-06 | 4.66E-08 | 0.024 | 0.019 | 0.054 |

Legend: *, p values were estimate using the hypergeometric distribution test; **, expected values were estimated based on the number of genes in the human genome (63,677) and the number of genes in the corresponding category of human-specific regulatory networks; ***, this category of genes was reported in Kronenberg et al. (2018); TE, transposable genetic elements; hESC, human embryonic stem cells; DLPFC, dorsolateral prefrontal cortex; MLME, multi lineage markers expression;
