## Supplemental Table S10 for "Analysis of genomic loci harboring 59,732 human-specific regulatory sequences reveals unique to human regulatory patterns associated with brain development"

**Supplemental Table S10.** Enrichment within human-specific regulatory networks of genes comprising expression signatures (GES) of human-specific transcriptional networks of induced pluripotent stem cells (iPSC).

**Networks of genes associated with expression of transposable elements (TE) in human dorsolateral prefrontal cortex**

| Classification category | Number of genes | hiPSC network | DOWN hiPSC vs NHP iPSC | UP hiPSC vs NHP iPSC |
| --- | --- | --- | --- | --- |
| Human genome | 63677 | 100 | 50 | 50 |
| Networks of genes associated with human DLPFC-expressed TE | 22863 | 76 | 40 | 36 |
| Percent | 35.90 | 76.00 | 80.00 | 72.00 |
| Enrichment** | 1.00 | 2.12 | 2.23 | 2.01 |
| P value* |  | 2.72E-16 | 1.90E-10 | 1.77E-07 |

**GES of the Multi-lineage Markers Expressing (MLME) cells of human preimplantation embryo**

| Classification category | Number of genes | hiPSC network | DOWN hiPSC vs NHP iPSC | UP hiPSC vs NHP iPSC |
| --- | --- | --- | --- | --- |
| Human genome | 63677 | 100 | 50 | 50 |
| GES of the MLME cells of human preimplantation embryo | 12735 | 52 | 22 | 30 |
| Percent | 20.00 | 52.00 | 44.00 | 60.00 |
| Enrichment** | 1.00 | 2.60 | 2.20 | 3.00 |
| P value* |  | 8.91E-13 | 7.15E-05 | 5.72E-10 |

**Regulatory networks of genes associated with human-specific structural variants*****

| Classification category | Number of genes | hiPSC network | DOWN hiPSC vs NHP iPSC | UP hiPSC vs NHP iPSC |
| --- | --- | --- | --- | --- |
| Human genome | 63677 | 100 | 50 | 50 |
| Genes associated with human-specific deletions and insertions | 10992 | 34 | 17 | 17 |
| Percent | 17.26 | 34.00 | 34.00 | 34.00 |
| Enrichment** | 1.00 | 1.97 | 1.97 | 1.97 |
| P value* |  | 2.44E-05 | 2.03E-03 | 2.03E-03 |

Legend: Legend: *, p values were estimate using the hypergeometric distribution test; **, expected values were estimated based on the number of genes in the human genome (63,677) and the number of genes in the corresponding category of human-specific regulatory networks; ***, this category of genes was reported in Kronenberg et al. (2018); TE, transposable genetic elements; hESC, human embryonic stem cells; DLPFC, dorsolateral prefrontal cortex; MLME, multi-lineage markers expression; NHP, non-human primates; iPSC, induced pluripotent stem cells; hiPSC, human iPSC;
